# Meta-metabolomic Responses of River Biofilms to Cobalt Exposure and Use of Dose-response Model Trends as an Indicator of Effects

**DOI:** 10.1101/2023.06.19.545533

**Authors:** Simon Colas, Benjamin Marie, Mathieu Milhe-Poutingon, Marie-Claire Lot, Amiel Boullemant, Claude Fortin, Séverine Le Faucheur

**Affiliations:** Universite de Pau et des Pays de l’Adour, E2S-UPPA, CNRS, IPREM, Pau, France; UMR 7245 CNRS/MNHN « Molécules de Communication et Adaptations des Micro-organismes », Muséum National d’Histoire Naturelle, Paris, France; TotalEnergies, Pole d’Études et de Recherche de Lacq, France; BCenv, Gardanne, France; Institut National de la Recherche Scientifique – Eau Terre Environnement, Québec, Canada

**Keywords:** Metals, Microbial ecotoxicology, Benchmark doses, Metabolites, Microcosms, CRIDeM, CRIDaM

## Abstract

Metabolites are low molecular-weight molecules produced during cellular metabolism. The global expression of the meta-metabolome (metabolomics at the community level) could thus potentially be used to characterize the exposure of an organism or a community to a specific stressor. Here, the meta-metabolomic fingerprints of mature biofilms were examined after 1, 3 and 7 days of exposure to five concentrations of cobalt (0, 1 x 10^-7^, 1 x 10^-6^, 5 x 10^-6^ and 1 x 10^-5^ M) in aquatic microcosms. The global changes in meta-metabolomic fingerprints were in good agreement with those of the other biological parameters studied (cobalt bioaccumulation, biomass, chlorophyll content). To better understand the dose-responses of the biofilm meta-metabolome, the untargeted LC-HRMS metabolomic data were further processed using the DRomics tool to build dose-response model curves and to calculate benchmark doses (BMD). These BMDs were aggregated into an empirical cumulative density function. A trend analysis of the metabolite dose-response curves suggests the presence of a concentration range inducing defense mechanisms (CRIDeM) between 4.7 x 10^-7^ and 2.7 x 10^-6^ M, and of a concentration range inducing damage mechanisms (CRIDaM) from 2.7 x 10^-6^ M to the highest Co concentration. The present study demonstrates that the molecular defense and damage mechanisms can be related to contaminant concentrations and represent a promising approach for environmental risk assessment of metals.

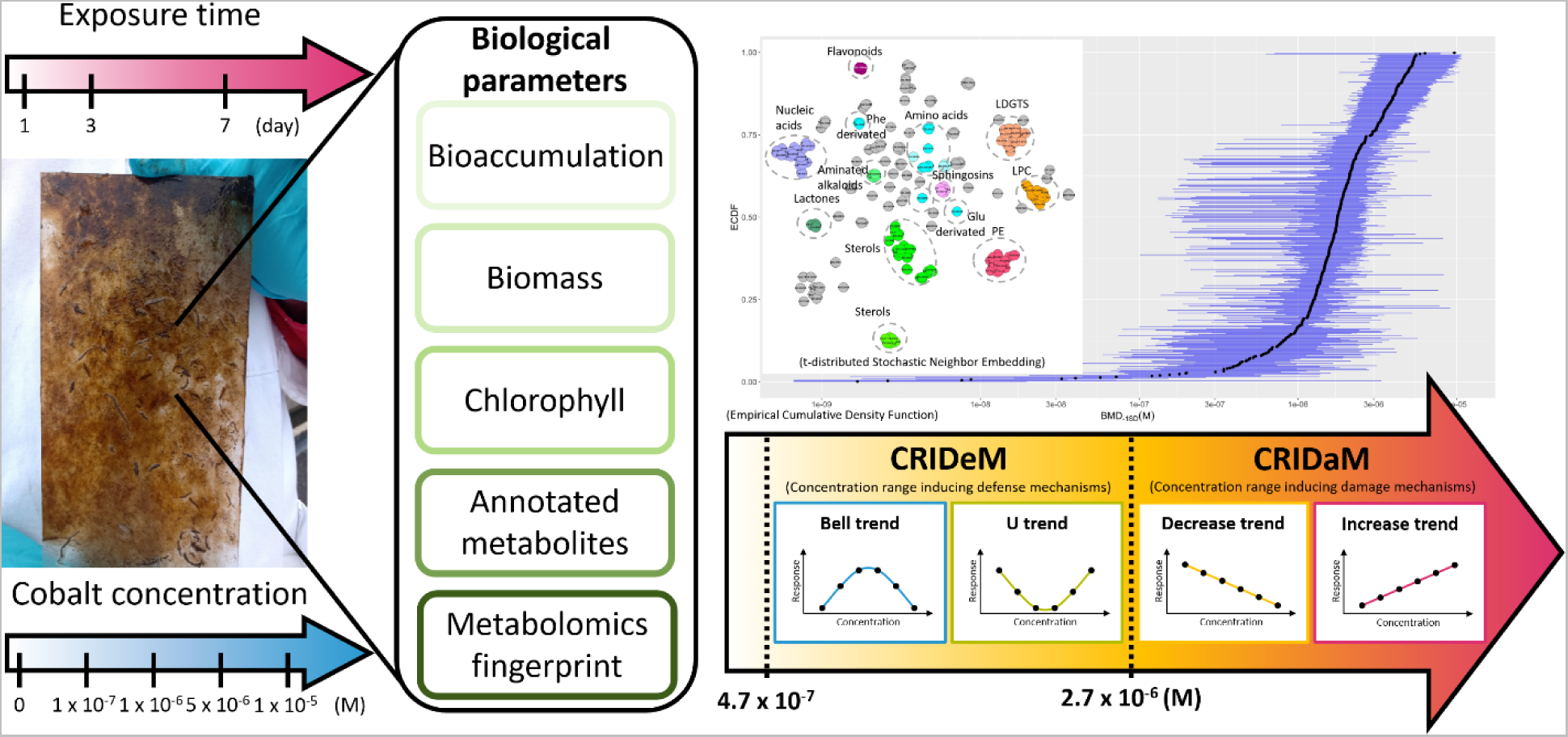

**SYNOPSIS:** This study focuses on the interpretation of the metabolite dose-response trends in river biofilms exposed to cobalt to identify concentration range inducing cellular mechanisms and improve the environmental risk assessment of metals.

## INTRODUCTION

In aquatic ecosystems, biofilms grow on submerged substrata and form the basis of the aquatic food web, as they include a large proportion of primary producers (algae and cyanobacteria) as well as bacteria, fungi and meiofauna. These communities are well known to accumulate metals as a function of their ambient concentration and speciation.^1–3^ Effects of metals on biofilms have been shown at all levels of the biological organization, from subcellular^4, 5^ and cellular^6–9^ levels to communities.^4, 5, 10^ Among these effects, modification of metabolite contents are linked with contaminant concentration and exposure to stressors.^11^

Metabolites are small molecular-weight molecules produced during metabolic processes. They have various cellular roles in basal functioning and defense mechanisms^11^ such as in the osmoregulation, in antioxidant activities, or metal chelation.^12^ They can also be related to damages caused by a stress such as lipid peroxidation^13^ or hypoxia.^14^

The whole metabolome is mainly examined using a non-targeted Liquid Chromatography-High Resolution Mass Spectrometry (LC-HRMS)^15^ and is often called the metabolomic fingerprint.^16^ In this approach, both unannotated and annotated metabolites are considered; the annotation consisting of assigning a tentative metabolite candidate on the basis of the mass of the ion, its retention time and its spectral fragmentation using a specific database. Metabolite responses are then often processed using multivariate statistical approach to distinguish between control and contaminant exposed group.^17^ Less frequently used, dose-response curves can also be drawn using the metabolite response, from which benchmark doses (BMD) – which correspond to the doses of the tested contaminants at which the response of the organisms differs from the control group – can be retrieved.^16, 18, 19^ These BMDs can be further aggregated in an empirical cumulative density function (ECDF) to get the whole metabolome response as a function of the contaminant concentrations.^16, 20^ Building metabolite ECDFs for biofilms exposed to contaminants such as metals may therefore provide valuable information on their exposure and effects.

Beside the building of ECDFs, dose-response curves for metabolomic response as a function of concentration might be further exploited with the analysis of their trends. Indeed, cellular defense mechanisms such as the activities of antioxidant enzymes^21, 22^, phytochelatins^23^ or biotransformation enzymes and associated compounds^23^ preferentially describe a bell-shaped or U-shaped trend. On the other hand, cellular damage mechanisms such as change in pigment contents^21, 22^, reactive oxygen species^21^ or malonaldehyde levels^24^ preferentially vary in a monotonically way (increasing or decreasing trend). In other words, the induction of a defense mechanism, the role of which is to have a positive impact on the organism, would begin following exposure to low-level stress, up to a certain threshold of stress intensity from which the organisms’ defense mechanisms are overwhelmed. The concentrations of the defense response would then decrease.^25^ This behaviour would explain a bell-shaped response. The U-shaped dose-response would follow the same biphasic response principle.^25^ For damage metabolites, their induction would mainly be initiated once the defense mechanisms have been overcome and would result from the continuous degradation of cellular and sub-cellular compounds, hence their tendency to increase and decrease in proportion to the intensity of the stress.^26^ Our working hypothesis is that the metabolome trend analysis coupled with the use of ECDF would provide information on the concentration range inducing defense mechanisms (CRIDeM) and the concentration range inducing damage mechanisms (CRIDaM), without the need of further identifying the metabolites. So far, no studies have used this approach for ecotoxicological studies.

The objective of the present study was thus to develop this innovative approach in biofilms exposed to cobalt (Co). Cobalt is an emerging contaminant due to its intensive use in the energy transition and has recently been identified as a relevant substance to be monitored in France^27^ and as a toxic substance in Canada^28^. Its effects on microorganism communities has been little studied, which limits its adequate risk assessment.^29–32^ Cobalt effect on biofilms will be examined using their meta-metabolome response as a function of Co concentrations and exposure time whereas the relevance of the calculated CRIDeM and CRIDaM will be assessed by comparing their range of induction with more traditional biological parameters (biomass and chlorophyll contents).

To that end, mature biofilms were exposed for 7 days to increasing concentrations of Co in microcosms placed in outdoor conditions. They were collected at three different step times (1, 3 and 7 days) to analyse Co bioaccumulation, biomass, chlorophyll content and meta-metabolome response. This present work provides significant evidences on the usefulness of the trend analysis of metabolite dose-response curves to evaluate the effects of a metal on communities.

## MATERIAL AND METHODS

### 1. Biofilm colonisation and exposure to Co in microcosms

Biofilm colonisation and exposure experiments were carried out at the outdoor TotalEnergies facility (pilot rivers) in Lacq (PERL, France). Four weeks before the start of the experiment, the mature biofilms were collected by natural colonisation of glass slides (5 x 10 cm). A total of 15 microcosms were filled with 15 L of the *Gave de Pau* water; three for each of the five exposure conditions: 2 x 10^-9^ M (background concentration used as control), 1 x 10^-7^, 1 x 10^-6^, 5 x 10^-6^ and 1 x10^-5^ M Co. Cobalt (Cobalt Standard for ICP, 1000 mg·L^-1^, Supelco, Germany) was added in each microcosm and let equilibrating for 24 h (D-1). The microcosms were placed in one of the artificial streams of the facility in order to maintain the water temperature at 13.5 ± 1.0°C (Table S2). Seven mature biofilm slides were placed in each microcosm (105 in total). After one, three and seven days of exposure (D1, D3 and D7), two biofilm slides were collected from each microcosm, one for Co accumulation and one for meta-metabolomics. At D7, one additional slide was sampled for the determination of chlorophyll content. The samples were stored in the dark at -20°C before analyses. The preparation of exposure media is further described in Text S1.

#### 1. 2. Water analysis and Co speciation

At D-1, D1, D3 and D7, 10 mL of exposure medium were taken from each microcosm in triplicates to determine metal concentrations. Samples for cation, anion and dissolved organic carbon (DOC) concentrations were collected at D-1. The sampling and analysis protocol of the different physico-chemical parameters of the exposure media are presented in Text S2.

#### 1. 3. Biofilm analysis

Biofilm samples collected for the determination of Co accumulation, chlorophyll content and meta-metabolome analyses were freeze-dried and weighed prior to further processing. These data were used to express the quantity of dry biomass (in g_DW_) per glass slide area (cm^2^). To obtain total and intracellular Co concentrations, the freeze-dried biofilms were digested with HNO_3_ 70% acid and H_2_O_2_ 30% before being mineralized (UltraWAVE™ oven, Milestone, Italy). To distinguish between total and intracellular Co content, biofilms were previously rinsed with 10 mM ethylenediaminetetraacetic (EDTA) for 10 min in order to remove adsorbed metals on the biofilm surface.^1,33^ The detailed protocol for the determination of bioaccumulated Co is provided in Text S3. Chlorophyll content was analysed by spectrophotometry and the quantification of the different pigments (chlorophyll *a*, chlorophyll *b* and chlorophylls *c*_1_ and *c*_2_) were determined following the Jeffrey and Humphrey protocol^34^. Further details are presented in Text S4. Meta-metabolomic analyses were carried out at the Muséum National d’Histoire Naturelle de Paris (MNHN) following the method of Le Moigne and his collaborators. ^35^ Briefly, freeze-dried biofilms (1 mg) were diluted in 10 µL of cold 75% methanol acidified with 0.1% formic acid and sonicated on ice during 30 s, at 80 % of the maximum intensity (SONICS Vibra Cell, Newton, CT, USA; 130 Watts, 20 kHz). The homogenates were then centrifuged at 4°C (12,000 g: 10 min). The supernatants were then collected and stored in the dark at -20°C. For the mass spectrometry analysis, 2 µL of the extracts were injected on an ultra-high-performance liquid chromatography (UH-PLC) (ELUTE, Bruker, Bremen, Germany) equipped with a Polar Advance II 2.5 pore C18 (Thermo Fisher Scientific, Waltham, MA, USA) chromatographic column. The molecule separation was obtained with a flow rate of 300 µL·min^-1^ under a linear gradient of acetonitrile (from 5 to 90% in 15 min) acidified with 0.1% formic acid. Metabolite contents were then analysed using an electrospray ionization hybrid quadrupole time-of-flight (ESI-QqTOF) high-resolution mass spectrometer (Compact, Brucker, Bremen, Germany) in the range of 50-1500 m/z. The global feature contents were extracted from raw data with Metaboscape 4.0 (Bruker) and annotation was attempted by molecular network approach performed with MetGem 1.3.6. The analysis method and annotation protocol are detailed in Text S5.

#### 1. 4. Dose-response models

Dose-response curves were built based on the biofilm meta-metabolomic response to Co^2+^ concentrations at D7 using the DRomics package under R software following the recommendations by EFSA Scientific Committee and Larras and his collaborators.^18, 19^ Pre-processing was performed on the raw data. First, the half-minimum method was applied for missing values. The data set was then log-2 transformed and the significantly responding metabolites were selected using an ANOVA with a false discovery rate (FDR) of 0.05. For each selected metabolite, a dose-response model was then constructed using the best AICc (second-order Akaike criterion) values as selected criteria. When dose-response curves could not be reliably fitted, the metabolite was removed from the analysis. The models were characterized according to the trends (bell-shaped, decreasing, increasing and U-shaped) of the fitted dose-response curves.

A benchmark-dose – 1SD (BMD_-1SD_) was calculated from the best model for each metabolite (Figure S1). This BMD_-1SD_ is the concentration corresponding to a benchmark response (BMR_-zSD_) defined as follows:

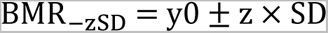

where y0 is the mean control response, SD is the residual standard deviation of the considered model, *i.e.*, 95% confidence interval (CI), and z is the factor of SD (z fixed at 1).^16^ When the calculated BMR_-1SD_ was within the range of response values defined by the model but the corresponding BMD_-1SD_ value was calculated to be outside the range of the tested Co concentrations, the metabolite was not taken into account (Figure S1). The CI on the BMD values were calculated by bootstrap (1000 iterations). After this step, models were removed when one of the bounds of their 95% CI could not be computed. Such a result could be obtained when bootstrapped BMD values were not reachable due to model asymptotes or to calculated values outside the range of tested doses. Finally, the distribution of all BMDs was compiled into an empirical cumulative density function (ECDF) to obtain an integrative response of biofilm exposure. Additional ECDFs were further constructed as a function of the dose-response curve trends.

#### 1. 5. Statistical treatments

The pseudo-replication due to the analysis of replicates within each tank was considered with the use of mixed models. To that end, statistical analyses were performed using the lme4 package of R software. The significance of Co exposure on Co bioaccumulation, biomass and chlorophyll content was assessed according to Dunn’s *post-hoc* test after Kruskal-Wallis non-parametric tests process on R software. Analyses of unannotated and annotated metabolites (MS analysis) were performed on MetaboAnalyst 5.0, including matrix normalization (Pareto), partial least square discriminant analysis (PLS-DA) and ANOVA (analysis of variance).

## RESULTS

### 1. Characterization of the Co content of the exposure medium

The *Gave de Pau* water used for exposure media preparation, had a mean pH of 7.90 ± 0.02, a background total dissolved concentration of 1.7 ± 0.2 x 10^-9^ M Co, 4.1 ± 0.1 x 10^-5^ M Ca, 2.9 ± 0.1 x 10^-6^ M Mg and 1.2 ± 0.3 mg DOC·L^-1^ (Table S1). Cobalt was predicted to be mainly under its free form Co^2+^ at a concentration of 1.1 ± 0.1 x 10^-9^ M, and the remaining species being mostly the carbonato-complex, CoCO_3_. A small percentage of the total Co (< 0.01 %) was calculated to be complexed by fulvic acid. The average total dissolved Co concentration, taking into account the three sampling days (D1, D3 and D7) for each nominal condition (exposure groups) 0, 1 x 10^-7^, 1 x 10^-6^, 5 x 10^-6^ and 1 x 10^-5^ M, were 3.1 ± 1.8 x 10^-9^, 1.4 ± 0.8 x 10^-7^, 1.6 ± 0.8 x 10^-6^, 8.6 ± 3.7 x 10^-6^ and 1.6 ± 0.7 x 10^-5^ M, respectively (Table S2). The corresponding calculated average Co^2+^ concentrations were 1.5 ± 0.9 x 10^-9^, 0.7 ± 0.6 x 10^-7^, 0.7 ± 0.5 x 10^-6^, 4.0 ± 0.9 x 10^-6^ and 0.8 ± 0.2 x 10^-5^ M, respectively (Table S2).

### 2. Metal bioaccumulation

Mature biofilms naturally contained 1.3 ± 0.1 x 10^-7^ mol·g_DW_^-1^ total accumulated Co (Table S3). Cobalt intracellular contents significantly increased with exposure concentrations (Figure 1A and Table S3) at each sampling time (D1, D3 and D7). After D7, the majority of accumulated Co by biofilms was found to be intracellular (70 ± 20 %; n=15). Background intracellular Co concentration was 1.23 ± 0.08 x 10^-7^ mol·g_DW_ ^-1^. Both total and intracellular Co accumulation were significantly correlated with total dissolved Co and Co^2+^ concentrations in the exposure media (FigureS2 and Table S3). The description of the obtained correlations between Co concentrations in exposure media and accumulated by biofilms are presented in Text S6. The accumulation of other metals (Li, Ni, Cu, Zn, Pb, As and Cd) naturally present in the exposure media was not impacted by the concentrations of Co in the exposure medium (Table S3).

**Figure 1:**
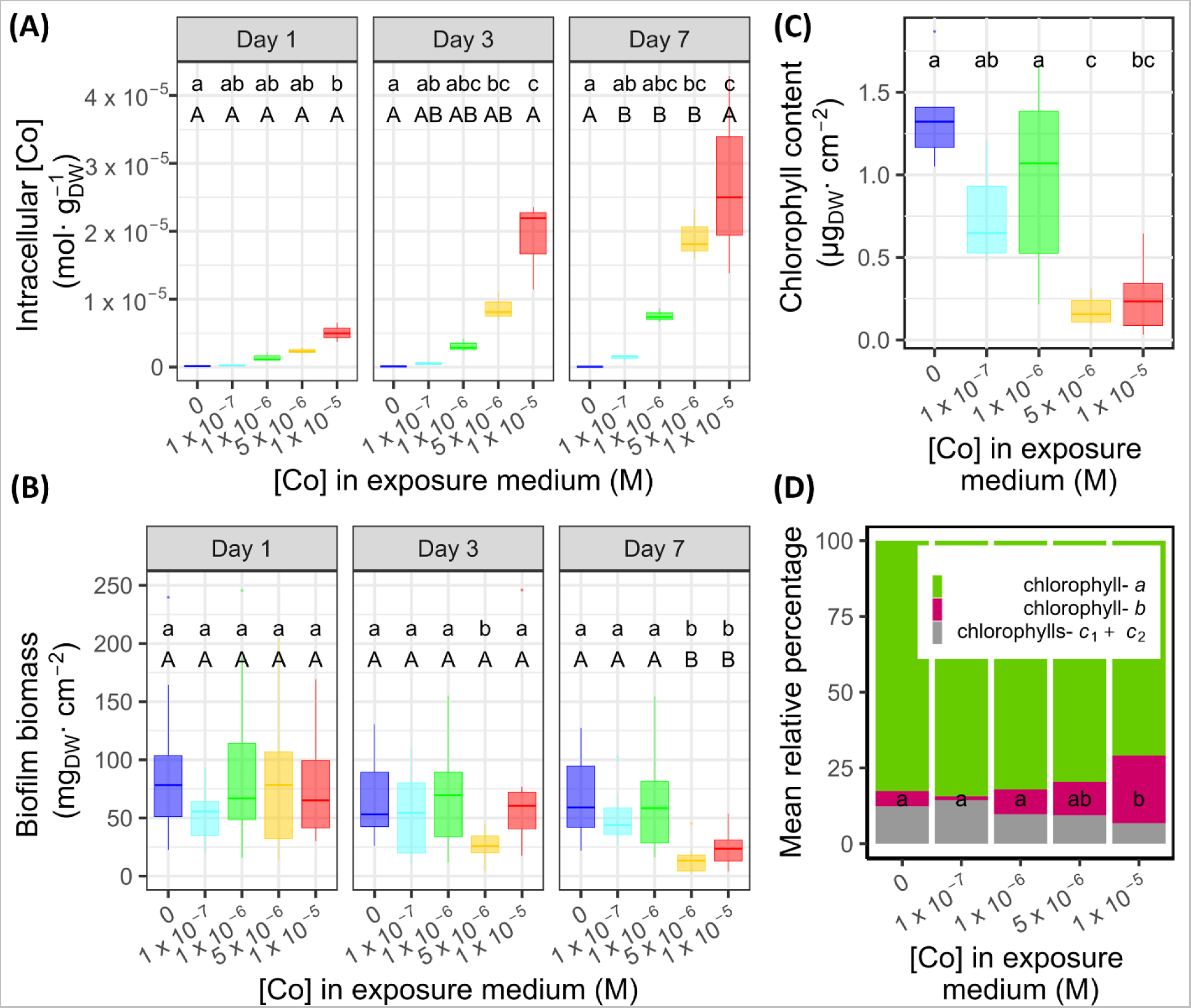
Accumulation and effects of Co on biomass and chlorophyll contents of river biofilms. (A) Intracellular Co content (mol·g^-1^_DW_) as function of total Co concentration in exposure medium (M) over time (day). (B) Biofilm biomass (mg_DW_·cm^-2^) as a function of time (day) according to exposure concentrations to Co (M). (C) Chlorophyll content (µg_DW_·cm^-2^) after 7 days of exposure according to exposure concentration Co (M). (D) Mean relative percentage of pigment composition in chlorophylls of biofilms exposed for 7 days to different Co concentrations. The lower-case letters correspond to the significative groups defined by a Dunn’s *post-hoc* test (*p*<0.05) performed after Kruskal-Wallis non-parametric tests among Co exposure concentrations at each exposure time and the capital letters correspond to the significant groups defined by a Dunn’s *post-hoc* test (p<0.05) performed after Kruskal-Wallis non-parametric tests among the exposure times for each exposure concentration.

### 3. Biomass

Biofilms exposed to natural background concentrations of Co and exposed to 1 x 10^-7^ and 1 x 10^-6^ M Co presented similar biomass throughout the whole duration of the experiment (Figure 1B). In contrast, their biomass significatively decreased over time when exposed to the highest concentrations (5 x 10^-6^ and 1 x 10^-5^ M). The greatest effect was observed at the concentrations of 5 x 10^-6^ and 1 x 10^-5^ M, concentrations at which the biomass decreased by 83% (from 80 to 13 mg_DW_·cm^-2^) and 69 % (77 to 24 mg_DW_·cm^-2^), respectively.

### 4. Chlorophyll content

Lesser content of the chlorophyll *a*, *b* and *c*_1_+*c*_2_ at D7 was observed as a function of Co concentration (Figure 1C). At the end of the experiment, biofilms exposed to natural Co concentrations had a total chlorophyll content of 0.73 µg_DW_·cm^-2^, while those exposed to the two highest concentrations had 0.10 and 0.15 µg_DW_·cm^-2^ of chlorophyll, respectively. After a week of exposure, the highest concentrations of Co (5 x 10^-6^ and 1 x 10^-5^ M) led to a lower chlorophyll content of up to 85%.

At D7, the relative composition of the different chlorophyll pigments was significatively different at the highest Co concentration compared to the control, 1 x 10^-7^ and 1 x 10^-6^ M (Figure 1D). Indeed, the relative contents of chlorophylls *a* and *c*_1_+*c*_2_ were not modified in biofilms exposed to 1 x10^-5^ M compare to the control, but their relative contents of chlorophyll *b* were 5.5-fold higher.

### 5. Meta-metabolomics response

In total, 2,117 metabolites were observed (Table S4) among all samples and 159 of them were annotated. Different families of molecules were attemptedly identified, including several lipids or lipid precursors (Lyso-diacylglyceryltrimethylhomoserine (LDGTS), sterol, phosphatidylethanolamine (PE), lysophosphatidylcholine (LPC), sphingosines) (Figures S3 and S5). The biofilm meta-metabolomic response was examined by comparing the samples at each exposure Co concentration and each exposure time, first for the whole 2,117 metabolites and second using the 159-annotated metabolite datasets.

**Figure 3:**
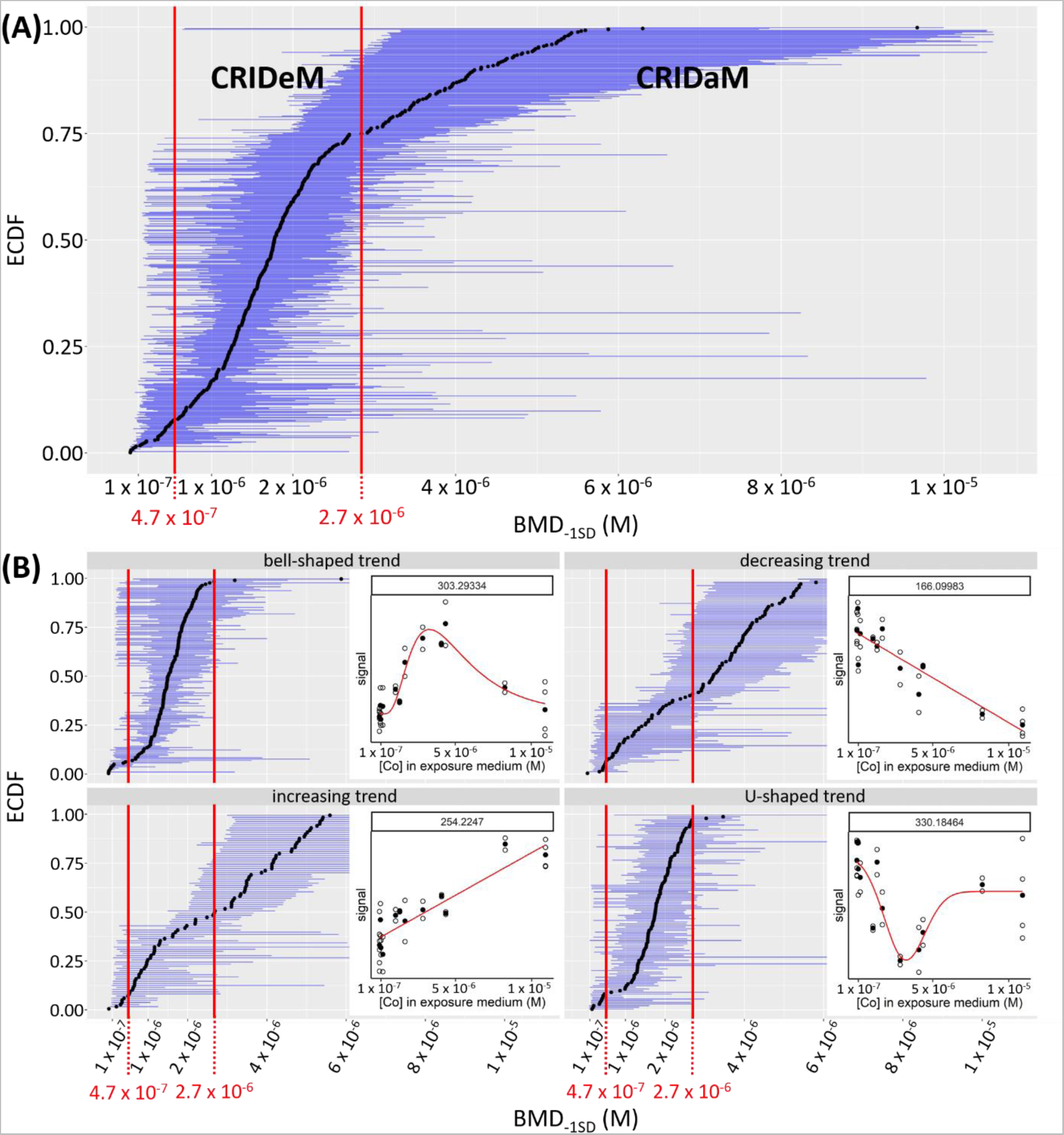
Meta-metabolomics dose-response models of biofilms after 7 days of exposure to Co. (A) Distribution of BMD-_1SD_ as an ECDF (Empirical Cumulative Density Function) with 95% CI of each BMD (M) in blue. (B) Distribution of BMD_-1SD_ as an ECDF split by trend of dose-response curves (bell-curved, decreasing, increasing and U-curved) with 95% CI of each BMD_-1SD_ (M) in blue. The red lines correspond to the concentrations delimiting the different concentration ranges inducing defense and damage mechanisms (CRIDeM and CRIDaM).

#### 5.1. Using all metabolites

Cobalt concentrations in the exposure media significantly impacted the levels of 489 metabolites and the exposure time affected 522 metabolites. The interaction of the two factors induced the modification of 502 metabolites (two-way ANOVA, p-value < 0.05). In total, out of 2,117 metabolites observed, 1,513 of them present relative content variations, which represents more than 71% of the whole meta-metabolome that could have been detected and analysed (Figure S. The effects of Co and exposure time on meta-metabolomic response were analysed using supervised partial least-squares discriminate analysis (PLS-DA). The constructed PLS-DA model showed significant R^2^ *cumulative*, Q^2^ *cumulative* (Table S5) and permutation score (0/2,000 permutations, p-value < 0.0005). The meta-metabolomic response of the control biofilms did not vary between D1 and D3 (Figure 2). However, at D7, a significant difference was observed compared to D1 and D3 (PERMANOVA, p-value < 0.05). At D1, the global meta-metabolomic signature was only different between biofilms exposed to 1 x 10^-7^ and 1 x 10^-5^ M Co (Figure 2B) (PERMANOVA, p-value < 0.05). At D3, the effect on the meta-metabolome was more important. Indeed, the meta-metabolomic signature between control biofilms was different from those of biofilms exposed to 5 x 10^-6^ and 1 x 10^-5^ M (PERMANOVA, p-values < 0.05). In addition, the signature of biofilms exposed to 1 x 10^-7^ M differed from that of biofilms exposed to 1 x 10^-5^ M, but not to those of biofilms exposed to the other studied concentrations (PERMANOVA, p-value < 0.05). The metabolic profile of biofilms exposed to 5 x 10^-6^ M was also different from biofilm exposed to 1 x 10^-5^ M (PERMANOVA, p-value < 0.01) (Figure 2C). At D7, the meta-metabolomic response was different between all the studied exposure concentrations, except for the control biofilms and those exposed to 1 x 10^-7^ M (PERMANOVA, p-values < 0.05) (Figure 2D). Finally, for each exposure condition, the meta-metabolomic signature of the biofilms was different between D1 and D7 (PERMANOVA, p-value < 0.05). The differences and the similarities in the biofilm meta-metabolomic profiles can also be observed on a heatmap with hierarchical classification representation (Euclidean distance, Figure 2E). The profiles of the control biofilms (at each time step), and those of the biofilms exposed to the lowest concentration at D1 gathered in the same group. The meta-metabolomic profiles of the biofilms exposed to the two highest concentrations at D7 were also grouped together.

**Figure 2:**
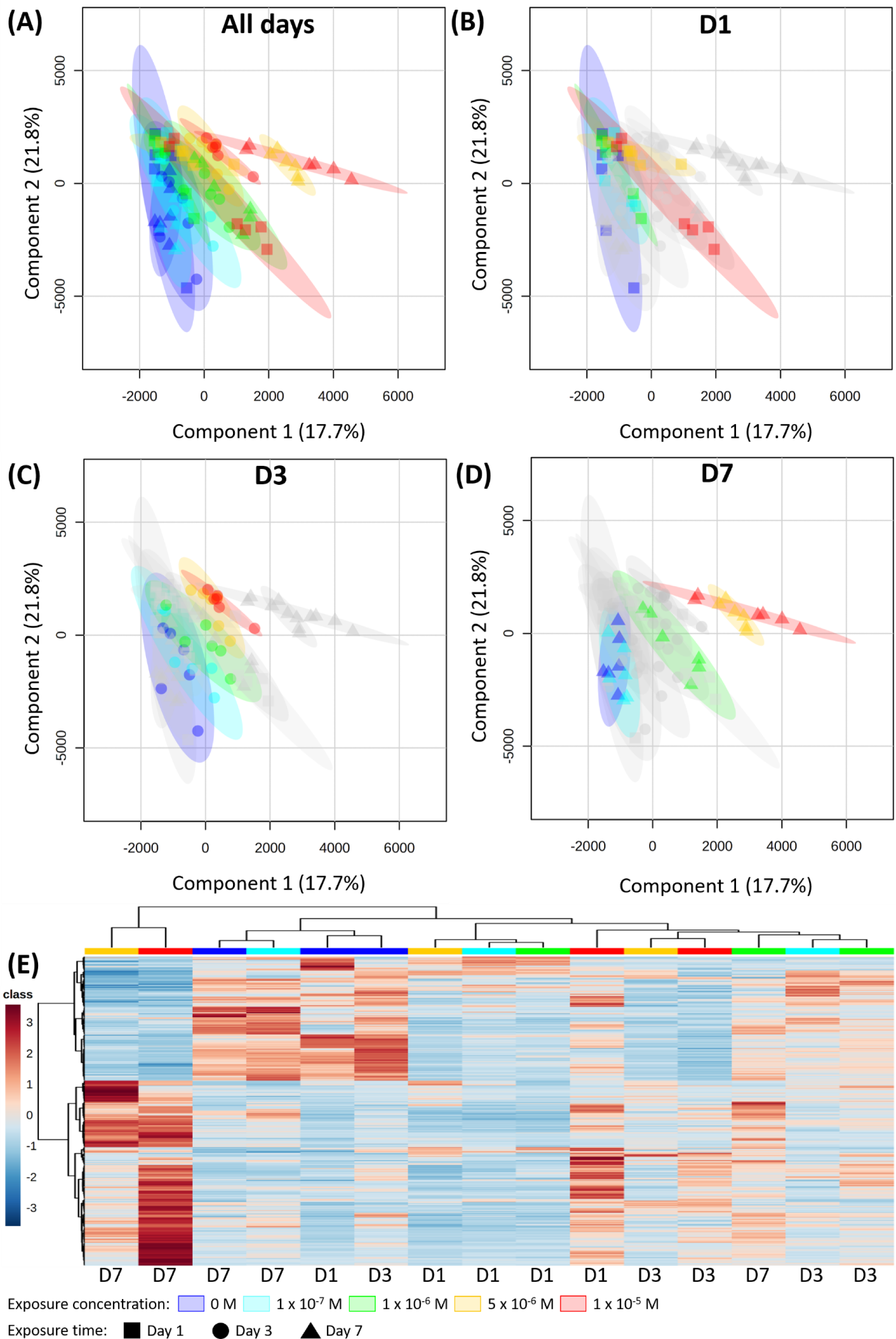
Meta-metabolomics response of biofilms according to time and concentrations of exposure to Co. (A) Individual score plot generated from PLS-DA analysis performed with 2,117 variables to components 1-2 with all conditions grouped together, (B) after one day of exposure, (C) three and (D) seven. (E) Four examples of representative box plot. (F) Heatmap with hierarchical classification representation of concentration class averages (Ward clustering according to Euclidian distances) performed from relative intensities of the dysregulated analytes with a VIP score > 1 for the component 1.

### 5.2. Using only annotated metabolites

Similar results were observed for the 159 annotated metabolites whose detailed analysis is presented in Text S7, Figures S3 and S5. Cobalt concentrations in the exposure media significantly induced a change in the production of 39 annotated metabolites. The exposure time disrupted the content of 25 annotated metabolites whereas the interaction of the two factors impacted the content of 33 metabolites (two-way ANOVA, p-value < 0.05). As such, out of 159 annotated metabolites, 61% of the biofilm metabolites were disturbed by the experimental treatments. A closer look at the meta-metabolomic profiles (Figures S3G and S3H) showed that exposure to Co caused greater production of sphingosines, PE and LDGTS as well as a decrease in the flavonoid content.

#### 5.3. Dose-response models

Dose-response curves were constructed to illustrate metabolite response as a function of Co concentrations at D7. Out of 2,117 metabolites, 892 metabolites were selected using an ANOVA with a FDR of 0.05. Using the AIC criteria to select the best fit models, 236 dose-response curves out of these 892 curves were removed as no model could be reliably fitted. From those modelled dose-response curves, three BMD_-1SD_ values could not be calculated as the BMR stands within the range of response values defined by the model but outside the range of tested doses. Also, bootstrap CI computation failed on eight metabolites due to the lack of convergence of the model fit for a fraction of the bootstrapped sampled greater than 0.5.

Benchmark doses (BMD_-1SD_) calculated from each fitted dose-response curve were grouped together via ECDF (Figure 3A and Table S6). One hundred and sixty-seven out of 656 BMD_-1SD_ values were removed because at least one bound of the 95% CI could not be computed due to bootstrapped BMD values which were not reachable. By comparing the proportions of metabolites impacted compared to the control within the range of tested concentrations, 2% of the metabolites were impacted between the control and 1 x 10^-7^ M Co, 15% were modified between 1 x 10^-7^ and 1 x 10^-6^ M whereas 78% of the metabolites had their response modified between 1 x 10^-6^ and 5 x 10^-6^ M. Finally, 5% of metabolites were impacted between these two highest concentrations.

The models were further classified into four different categories as a function of their respective curve trend (Figure 3B and Table S6). As such, 151 exhibited a bell-shaped curve, 92 an increasing trend, 121 a decreasing trend and 117 a U-shaped curve (Figure 3B). Using the ECDF curve, three main concentration ranges of meta-metabolomic response could be distinguished: the first one ranging from 0 to 4.7 x 10^-7^ M, the second from 4.7 x 10^-7^ M to 2.7 x 10^-6^ M and the third one from 2.7 x 10^-6^ M to the highest concentration (Figure 3B). In the first concentration range, 6% of metabolites describing a bell-shaped response curve were present, 92% in the second one and 1% in the third range. Eight % of the metabolites with a U-shaped response curve were present in the first concentration range, 89% in the second one and 3% in the third range. Seven % of the metabolites with an increasing response curve were present in the first concentration range, 43% in the second one and 50% in the third range. Finally, 4% of the metabolites with a decreasing response curve were present in the first concentration range, 36% in the second one and 60% in the third range. (Figures 3A, 3B and Table S6). The proportion of metabolites with a bell-shaped and U-shaped dose-response trend was significantly greater in the second concentration range than in the third concentration range (Chi2 = 182.28, d.f. = 6, p-value < 0.001). Conversely, the proportion of metabolites with increasing and decreasing dose-response trends was significantly greater in the third range than in the second range (Chi2 = 182.28, d.f. = 6, p-value < 0.001).

## DISCUSSION

### 1. Biofilm meta-metabolomic response as a function of Co concentration and exposure time

As expected, Co accumulation increased linearly as a function of the calculated free ion Co^2+^ concentration (Figure S1). Published data on Cu, Zn, Cd and Ni accumulation in biofilms obtained either in the laboratory or in the field have shown similar correlations.^1–3, 36^ That metal accumulation is the resultant of metal uptake and release processes, explaining the very dynamic nature of bioaccumulation by biofilms and its tight relationship with the ambient metal concentrations.^1^ This relationship between Co accumulation and ambient concentration was already visible after 1 day. Bioaccumulation is the first step before the induction of an effect in organisms. However, although bioaccumulation was significantly different between the highest exposure Co concentrations and, control and 1 x 10^-7^ M at D1, no statistical modification was found at the meta-metabolome level between control and Co-exposed biofilms, except between 1 x 10^-7^ and 1 x 10^-5^ M (Figure 1A and 2B). Metabolite change has been reported to occur within minutes after contaminant exposure, when analysing specific biomolecules one at a time and involved in detoxification and damage processes.^37, 38^ For example, microalgae exposed to Cu, Cd and Pb enzymatically produced metal-binding phytochelatins within the first 15 min of exposure.^39, 40^ Using targeted metabolomic fingerprints, microalgae exposed to Ag nanoparticles (97 metabolites studied) ^41^ and Hg (93 metabolites studied) ^42^ were shown to respond within 2 hours. Changes in untargeted meta-metabolome (2,117 metabolites studied in this experiment) over time in a whole community of microorganisms such as freshwater biofilms had rarely been examined so far. A set of metabolites inherently describes a greater diversity in term of toxicological response typology and also range of intensity than a single metabolite. Measuring a significant effect implies that i) a large number of metabolites are affected and/or ii) the effect is greater than the control. Despite these challenges, a recent study successfully demonstrated a shift in biofilm meta-metabolome after only 15 min of exposure to 5 and 50 μg·L^-1^ diuron, an herbicide inhibiting the algal photosystem II. ^16^ The shift was shown to be concentration-dependent, with a higher number of metabolites affected at higher concentrations. The lack of a significant difference in the meta-metabolome response at D1 in our study might therefore be due to an exposure concentration to an essential metal that was too low to trigger a significant response in that short time step. Statistical differences were, however, measured at D3 between the control and the two highest exposure concentrations. This is also the case at D7 between almost all treatments. Indeed, at that exposure time, a clear pattern was observed with an effect on meta-metabolome intensifying with increasing Co concentrations. All these responses were statistically different, except for the control and 1 x 10^-7^ M. The change in the meta-metabolomic fingerprint of the biofilms depends thus both on the concentrations tested and the exposure time. This phenomenon can either be due to specific cellular responses at the organism level and/or a change in the microorganism community composition.

The use of the annotated metabolites provided similar results as a function of Co concentration and exposure time as those observed in the whole meta-metabolome analysis. Nevertheless, the latter provided insights on the cellular mechanisms modified by Co. The most noticeable effect observed on the annotated metabolites was the modification of the lipid contents (Figures S3G and S3H). Specifically, increased levels of sphingosines, PE, and LDGTS, which are lipids or lipid precursors, were observed. Several processes may explain these modifications: the formation of vacuoles made of lipid bilayers for metal sequestration,^43^ the repair of potential membrane damages caused by indirect oxidative stress,^44^ and the need for an additional internal source of energy in order to offset cellular and subcellular damages.^45^ Finally, modifications of lipid profiles may also be due to a shift in microorganism community, as the metabolome variation over time and the changes in pigment compositions could suggest (Figure 1D).^46^ Recent studies combining the analysis of metagenomics and meta-metabolomics have demonstrated a change in the lipid profiles and taxa composition of stressed biofilms exposed to erythromycin and Ag nanoparticles, although the biofilm microbial diversity was not altered.^15, 47^ Overall, our results add up to those highlighting the modification of the biofilm lipid contents when exposed to contaminants.^15, 16, 48^

### 2. Comparison of meta-metabolomic response using the BMD approach with the measured biological parameters

Cobalt negatively impacted biofilm biomass and chlorophyll content (Figures 1B and 1C). This effect has already been observed in biofilms exposed to other metals such as Cd^6^, Cu^49^ and Zn.^50^ Because of their ability to bind to important biomolecules, metals can affect the metabolism of algae and cyanobacteria, in particular their photosynthesis mechanisms.^51, 52^ Cobalt has an adverse effect on the relative proportions of the different pigments within the different exposure groups at D7, with an increase in chlorophyll *b* proportion at the highest concentration (Figure 1D). This may reflect a shift in the biofilm community. Indeed, in an aquatic environment, chlorophyll *b* indicates the presence of green algae (chlorophyceae, prasinophyceae, euglenophyceae)^53–55^ and chlorophylls *c* indicates so-called brown algae, notably diatoms. ^53, 54, 56, 57^ The high concentrations of Co would therefore lead to an increased presence of green algae within the communities forming the biofilms.

Within the range of tested Co concentrations, the use of BMDs highlighted that 78 % of the metabolites had their response modified between 1 x 10^-6^ and 5 x 10^-6^ M (Figure 3A). Effects on chlorophyll pigments and biomass were in good agreement with this result. Indeed, Co effect was also observed to intensify between 1 x 10^-6^ and 5 x 10^-6^ M for both biological parameters (Figure 1B and 1C).

The approach for detecting the effects of stressor via the use of BMDs was first used in toxicology to address the limits of predictors such as the effective concentration (EC_x_), NOAEL or the lowest-observed adverse effect level (LOAEL) and to harmonise these predictive models of critical dose. ^58, 59^ Indeed, the use of BMDs possess several advantages ^60–62^, *e.g.* they (i) take into account the uncertainty related to the experiment and the inter-individual variability; (ii) consider the entire dose-response relationship; (iii) have a high precautionary principle ; and (iv) associate a risk with each dose tested. As such, the BMDs approach is also gaining interest in ecotoxicology. ^63^ In the present study, the results observed via the use of BMDs (Figure 3A) are in good agreement with the observed and quantifiable biological parameters (biomass and chlorophyll content) despite being retrieved from predictive models (Figures 1B and 1C).

Stubblefield and his collaborators determined the values of median hazardous concentration (HC) of 5% (HC_5,50%_) based on a species sensitivity distribution for freshwater organisms to be 3.1 x 10^-8^ M for chronic toxicity using EC_10_ values and 1.2 x 10^-7^ M using EC_20_ values.^64^ In our study, for both values, no effect on the biomass and the chlorophyll contents were observed in biofilms (Figures 1B, 1C and 1D) and the biofilm meta-metabolomic fingerprints were similar to those of the control at 10^-7^ M (Figures 2 and S3). The first observable effect concentration threshold on the ECDF was at 4.6 x 10^-7^ M Co, suggesting that our results are in line with those of Stubblefield *et al*.

### 3. Using trends of dose-response curves to identify biofilm response to Co

The analysis of Co effect on biofilm meta-metabolome via the calculation of BMDs and the use of ECDF made it possible to distinguish three distinct Co concentration ranges according to the meta-metabolomic dose-response (Figure 3B). The range 1 (0 - 4.6 x 10^-7^ M; 0 being the control) would correspond to effects induced by low Co concentrations in the ambient water. Since Co is a trace element, these meta-metabolomic changes might represent homeostasis processes,^65–68^ such as the beneficial uptake of Co required for microalgae optimum growth. ^69^ The range 2 (4.6 x 10^-7^ to 2.7 x 10^-6^ M) comprised mainly of metabolites describing bell- or U-shaped dose-response models. Both bell- or U-shaped trends correspond to those of well-known defense biomarkers such as antioxidant enzymes.^21, 70–72^ For example, in microalgae, catalase and superoxide dismutase activities were found to respond in a bell-shape manner with Cd concentrations.^21^ In this range 2, the microorganisms would set up their defense mechanisms in order to counter the increase in intracellular Co content and its potential harmful effects (Figure 1A and S1). We have defined this process as CRIDeM. The range 3 (concentrations above 2.7 x 10^-6^ M), the metabolite presenting significant response exhibited mainly increasing or decreasing trends. These trends would thus describe the microorganism cellular state when defense mechanisms are overwhelmed and molecular or cellular damages are occurring. In that range, damage biomarkers are commonly observed. ^21, 73–76^ For example, malondialdehyde contents resulting from lipid peroxidation increase proportionally to Cu, Zn and Cd concentrations in microalgae. ^24, 77^ Conversely, the contents of photosynthetic pigments decreased proportionally with Cr and Cd concentrations. ^77, 78^ We have defined this process as CRIDaM. For Co, the identified threshold value of 2.7 x 10^-6^ M, which might separate CRIDeM and CRIDaM, is in line with the changes observed using the traditional biological parameters studied here. Indeed, biofilm biomass decreased slightly at 1 x 10^-6^ M and much more significantly at 5 x 10^-6^ and 1 x 10^-5^ M (Figure 1B). The effect of Co on the chlorophyll content strongly changed between 1 x 10^-6^ and 5 x 10^-6^ M (Figure 1C). This CRIDaM threshold of 2.7 x 10^-6^ M is in good agreement with studies highlighting the harmful effects of Co on microalgae at the µM level. ^52, 79^

The complementary use of the construction of dose-response models, calculation of BMDs and trend analysis of the model curves were demonstrated to be a promising approach to interpret non-targeted metabolomics. Indeed, the response of microorganism communities to stress could be evaluated, without the limitation of metabolite annotation. Moreover, the interpretation of trends of metabolite dose-response models made it possible to identify and differentiate a CRIDeM and a CRIDaM, providing information on biofilm stress level. We have chosen to give these general terms of concentration ranges inducing defense or damage mechanisms and be not specific to metabolites in order to leave open the possibility of testing this concept with other omics studies (transcriptomics, proteomics, etc.). It would also be possible to test this interpretation with the effect of exposure time and verify whether the time range inducing defense mechanisms (TRIDeM) and time range inducing damage mechanisms (TRIDaM) concept would also be applicable. This technique of analysis and data processing is a promising avenue to be implemented in the risk assessment of contaminants in ecosystems.

## ASSOCIATED CONTENT

Description of the biofilm colonization and the Co exposure in microcosms, the method of the water analysis, Co speciation, total and intracellular Co concentration, chlorophyll content and meta-metabolomics analysis. And the detailed results of the physico-chemical parameters of the *Gave de Pau* water and the microcosms, the metal bioaccumulation, the effect of Co on annotated metabolites and the different parameters of the ECDF.

## AUTHOR INFORMATION

### Author Contributions

Conceptualization, S.C. and S.L F.; data collection, S.C., M.M-P. and S.L F.; water analysis, M.M-P.; bioaccumulation analysis; S.C., M.M-P.; biomass analysis, S.C.; chlorophyll analysis, S.C.; meta-metabolomic analysis, S.C. and B.M.; data processing, S.C., B.M. and S.L F.; writing-original draft preparation, S.C. and S.L F.; writing-review and editing, S.C., B.M., M.M-P., M-C.L., A.B., C.F. and S.L F.; funding acquisition, S.L F.. All authors have given approval to the final version of the manuscript.

### Notes

The authors declare no competing financial interest.

## ACKNOWLEDGMENT

We would like to thank the members of the Pôle Environment & Développement Durable, Pôle d’Etudes et de Recherche de Lacq (PERL) for access to the TotalEnergies facilities for microcosms exposure and the UMR 7245 MCAM, Muséum National d’Histoire Naturelle, Paris, France, especially the “Cyanobactéries, Cyanotoxines et Environnement” (CCE) team and the Plateau technique de spectrométrie de masse bio-organique (PtSMB) for laboratories and technical facilities for meta-metabolomics analysis. We also thank Patrick Baldoni-Andrey (TotalEnergies) and Nick Gurieff (Rio Tinto) for the scientific discussions.

## ABBREVIATIONS

BMD: benchmark dose
BMR: benchmark response
CRIDaM: concentration range inducing damage mechanisms
CRIDeM: concentration range inducing defense mechanisms
DOC: dissolved organic carbon
EC_x_: effective concentration
ECDF: empirical cumulative density function
HC: hazardous concentration
LDGTS: lyso-diacylglyceryltrimethylhomoserine
LOAEL: low-observed adverse effect level
LPC: lysophosphatidylcholine
NOAEL: non-observed adverse effect level
PE: phosphatidylethanolamine
TRIDaM: time range concentration inducing damage mechanisms
TRIDeM: time range concentration inducing defense mechanisms.

## REFERENCES

(1) Meylan, S.; Behra, R.; Sigg, L. Accumulation of Copper and Zinc in Periphyton in Response to Dynamic Variations of Metal Speciation in Freshwater. Environ. Sci. Technol. 2003, 37 (22), 5204–5212. https://doi.org/10.1021/es034566+.

(2) Lavoie, I.; Lavoie, M.; Fortin, C. A Mine of Information: Benthic Algal Communities as Biomonitors of Metal Contamination from Abandoned Tailings. Sci. Total Environ. 2012, 425, 231–241. https://doi.org/10.1016/j.scitotenv.2012.02.057.

(3) Laderriere, V.; Paris, L.-E.; Fortin, C. Proton Competition and Free Ion Activities Drive Cadmium, Copper, and Nickel Accumulation in River Biofilms in a Nordic Ecosystem. Environments 2020, 7 (12), 112. https://doi.org/10.3390/environments7120112.

(4) Barranguet, C.; Greijdanus, M.; Sinke, J. J.; Admiraal, W. Copper-Induced Modifications of the Modifications of the Trophic Relations in Riverine Algal-Bacterial Biofilms. Environ. Toxicol. Chem. 2003, 22 (6), 1340–1349. https://doi.org/10.1002/etc.5620220622.

(5) Bonet, B.; Corcoll, N.; Acuňa, V.; Sigg, L.; Behra, R.; Guasch, H. Seasonal Changes in Antioxidant Enzyme Activities of Freshwater Biofilms in a Metal Polluted Mediterranean Stream. Sci. Total Environ. 2013, 444, 60–72. https://doi.org/10.1016/j.scitotenv.2012.11.036.

(6) Duong, T. T.; Morin, S.; Coste, M.; Herlory, O.; Feurtet-Mazel, A.; Boudou, A. Experimental Toxicity and Bioaccumulation of Cadmium in Freshwater Periphytic Diatoms in Relation with Biofilm Maturity. Sci. Total Environ. 2010, 408 (3), 552–562. https://doi.org/10.1016/j.scitotenv.2009.10.015.

(7) Lavoie, M.; Fortin, C.; Campbell, P. G. C. Influence of Essential Elements on Cadmium Uptake and Toxicity in a Unicellular Green Alga: The Protective Effect of Trace Zinc and Cobalt Concentrations. Environ. Toxicol. Chem. 2012, 31 (7), 1445–1452. https://doi.org/10.1002/etc.1855.

(8) Morin, S.; Duong, T. T.; Herlory, O.; Feurtet-Mazel, A.; Coste, M. Cadmium Toxicity and Bioaccumulation in Freshwater Biofilms. Arch. Environ. Contam. Toxicol. 2008, 54 (2), 173–186. https://doi.org/10.1007/s00244-007-9022-4.

(9) Morin, S.; Lambert, A. S.; Rodriguez, E. P.; Dabrin, A.; Coquery, M.; Pesce, S. Changes in Copper Toxicity towards Diatom Communities with Experimental Warming. J. Hazard. Mater. 2017, 334, 223–232. https://doi.org/10.1016/j.jhazmat.2017.04.016.

(10) Doose, C.; Fadhlaoui, M.; Morin, S.; Fortin, C. Thorium Exposure Drives Fatty Acid and Metal Transfer from Biofilms to the Grazer *Lymnaea* Sp. Environ. Toxicol. Chem. 2021, 40 (8), 2220–2228. https://doi.org/10.1002/etc.5067.

(11) Gauthier, L.; Tison-Rosebery, J.; Morin, S.; Mazzella, N. Metabolome Response to Anthropogenic Contamination on Microalgae: A Review. Metabolomics 2020, 16 (1), 8. https://doi.org/10.1007/s11306-019-1628-9.

(12) Arora, N.; Dubey, D.; Sharma, M.; Patel, A.; Guleria, A.; Pruthi, P. A.; Kumar, D.; Pruthi, V.; Poluri, K. M. NMR-Based Metabolomic Approach To Elucidate the Differential Cellular Responses during Mitigation of Arsenic(III, V) in a Green Microalga. ACS Omega 2018, 3 (9), 11847–11856. https://doi.org/10.1021/acsomega.8b01692.

(13) Rocchetta, I.; Mazzuca, M.; Conforti, V.; Ruiz, L.; Balzaretti, V.; de Molina, M. del C. R. Effect of Chromium on the Fatty Acid Composition of Two Strains of Euglena Gracilis. Environ. Pollut. 2006, 141 (2), 353–358. https://doi.org/10.1016/j.envpol.2005.08.035.

(14) Zhang, W.; Tan, N. G. J.; Li, S. F. Y. NMR-Based Metabolomics and LC-MS/MS Quantification Reveal Metal-Specific Tolerance and Redox Homeostasis in Chlorella Vulgaris. Mol. Biosyst. 2014, 10 (1), 149–160. https://doi.org/10.1039/c3mb70425d.

(15) Pu, Y.; Pan, J.; Yao, Y.; Ngan, W. Y.; Yang, Y.; Li, M.; Habimana, O. Ecotoxicological Effects of Erythromycin on a Multispecies Biofilm Model, Revealed by Metagenomic and Metabolomic Approaches. Environ. Pollut. 2021, 276, 116737. https://doi.org/10.1016/j.envpol.2021.116737.

(16) Creusot, N.; Chaumet, B.; Eon, M.; Mazzella, N.; Moreira, A.; Morin, S. Metabolomics Insight into the Influence of Environmental Factors in Responses of Freshwater Biofilms to the Model Herbicide Diuron. Environ. Sci. Pollut. Res. 2022, 29 (20), 29332–29347. https://doi.org/10.1007/s11356-021-17072-7.

(17) Godzien, J.; Gil de la Fuente, A.; Otero, A.; Barbas, C. Metabolite Annotation and Identification. In Comprehensive Analytical Chemistry; Elsevier, 2018; Vol. 82, pp 415–445. https://doi.org/10.1016/bs.coac.2018.07.004.

(18) Larras, F.; Billoir, E.; Baillard, V.; Siberchicot, A.; Scholz, S.; Wubet, T.; Tarkka, M.; Schmitt-Jansen, M.; Delignette-Muller, M.-L. DRomics: A Turnkey Tool to Support the Use of the Dose–Response Framework for Omics Data in Ecological Risk Assessment. Environ. Sci. Technol. 2018, 52 (24), 14461–14468. https://doi.org/10.1021/acs.est.8b04752.

(19) EFSA Scientific Committee; Hardy, A.; Benford, D.; Halldorsson, T.; Jeger, M. J.; Knutsen, K. H.; More, S.; Mortensen, A.; Naegeli, H.; Noteborn, H.; Ockleford, C.; Ricci, A.; Rychen, G.; Silano, V.; Solecki, R.; Turck, D.; Aerts, M.; Bodin, L.; Davis, A.; Edler, L.; Gundert-Remy, U.; Sand, S.; Slob, W.; Bottex, B.; Abrahantes, J. C.; Marques, D. C.; Kass, G.; Schlatter, J. R. Update: Use of the Benchmark Dose Approach in Risk Assessment. EFSA J. 2017, 15 (1). https://doi.org/10.2903/j.efsa.2017.4658.

(20) Larras, F.; Billoir, E.; Scholz, S.; Tarkka, M.; Wubet, T.; Delignette-Muller, M.-L.; Schmitt-Jansen, M. A Multi-Omics Concentration-Response Framework Uncovers Novel Understanding of Triclosan Effects in the Chlorophyte Scenedesmus Vacuolatus. J. Hazard. Mater. 2020, 397, 122727. https://doi.org/10.1016/j.jhazmat.2020.122727.

(21) Cheng, J.; Qiu, H.; Chang, Z.; Jiang, Z.; Yin, W. The Effect of Cadmium on the Growth and Antioxidant Response for Freshwater Algae Chlorella Vulgaris. SpringerPlus 2016, 5 (1), 1290. https://doi.org/10.1186/s40064-016-2963-1.

(22) Ran, X.; Liu, R.; Xu, S.; Bai, F.; Xu, J.; Yang, Y.; Shi, J.; Wu, Z. Assessment of Growth Rate, Chlorophyll a Fluorescence, Lipid Peroxidation and Antioxidant Enzyme Activity in Aphanizomenon Flos-Aquae, Pediastrum Simplex and Synedra Acus Exposed to Cadmium. Ecotoxicology 2015, 24 (2), 468–477. https://doi.org/10.1007/s10646-014-1395-3.

(23) Navarrete, A.; González, A.; Gómez, M.; Contreras, R. A.; Díaz, P.; Lobos, G.; Brown, M. T.; Sáez, C. A.; Moenne, A. Copper Excess Detoxification Is Mediated by a Coordinated and Complementary Induction of Glutathione, Phytochelatins and Metallothioneins in the Green Seaweed Ulva Compressa. Plant Physiol. Biochem. 2019, 135, 423–431. https://doi.org/10.1016/j.plaphy.2018.11.019.

(24) Li, M.; Hu, C.; Zhu, Q.; Chen, L.; Kong, Z.; Liu, Z. Copper and Zinc Induction of Lipid Peroxidation and Effects on Antioxidant Enzyme Activities in the Microalga Pavlova Viridis (Prymnesiophyceae). Chemosphere 2006, 62 (4), 565–572. https://doi.org/10.1016/j.chemosphere.2005.06.029.

(25) Davis, J. M.; Svendsgaard, D. J. U-Shaped Dose-response Curves: Their Occurrence and Implications for Risk Assessment. J. Toxicol. Environ. Health 1990, 30 (2), 71–83. https://doi.org/10.1080/15287399009531412.

(26) Swenberg, J. A.; Fryar-Tita, E.; Jeong, Y.-C.; Boysen, G.; Starr, T.; Walker, V. E.; Albertini, R. J. Biomarkers in Toxicology and Risk Assessment: Informing Critical Dose– Response Relationships. Chem. Res. Toxicol. 2008, 21 (1), 253–265. https://doi.org/10.1021/tx700408t.

(27) Institut national de l’environnement industriel et des risques. Substances Pertinentes à Surveiller (SPAS) Dans Les Eaux de Surface - Etude Des Données à l’échelle Des Bassins Hydrographiques et Selon Les Types de Pression Chimique; Ineris-203220-2733069-v2.0; Ineris: Verneuil-en-Halatte, 2022. https://www.ineris.fr/fr/substances-pertinentes-surveiller-spas-eaux-surface-etude-donnees-echelle-bassins-hydrographiques (accessed 2023-06-02).

(28) Canada Gazette. *Order Adding a Toxic Substance to Schedule 1 to the Canadian Environmental Protection Act*, 1999: SOR/2019-197; Vol. 153. https://gazette.gc.ca/rp-pr/p2/2019/2019-06-26/html/sor-dors197-eng.html (accessed 2023-06-02).

(29) Barrio-Parra, F.; Elío, J.; De Miguel, E.; García-González, J. E.; Izquierdo, M.; Álvarez, R. Environmental Risk Assessment of Cobalt and Manganese from Industrial Sources in an Estuarine System. Environ. Geochem. Health 2018, 40 (2), 737–748. https://doi.org/10.1007/s10653-017-0020-9.

(30) Bhutiani, R.; Kulkarni, D. B.; Khanna, D. R.; Gautam, A. Geochemical Distribution and Environmental Risk Assessment of Heavy Metals in Groundwater of an Industrial Area and Its Surroundings, Haridwar, India. Energy Ecol. Environ. 2017, 2 (2), 155–167. https://doi.org/10.1007/s40974-016-0019-6.

(31) Khan, Z. I.; Arshad, N.; Ahmad, K.; Nadeem, M.; Ashfaq, A.; Wajid, K.; Bashir, H.; Munir, M.; Huma, B.; Memoona, H.; Sana, M.; Nawaz, K.; Sher, M.; Abbas, T.; Ugulu, I. Toxicological Potential of Cobalt in Forage for Ruminants Grown in Polluted Soil: A Health Risk Assessment from Trace Metal Pollution for Livestock. Environ. Sci. Pollut. Res. 2019, 26 (15), 15381–15389. https://doi.org/10.1007/s11356-019-04959-9.

(32) Xu, F.; Wang, Y.; Chen, X.; Liang, L.; Zhang, Y.; Zhang, F.; Zhang, T. Assessing the Environmental Risk and Mobility of Cobalt in Sediment near Nonferrous Metal Mines with Risk Assessment Indexes and the Diffusive Gradients in Thin Films (DGT) Technique. Environ. Res. 2022, 212, 113456. https://doi.org/10.1016/j.envres.2022.113456.

(33) Olguín, E. J.; Sánchez-Galván, G. Heavy Metal Removal in Phytofiltration and Phycoremediation: The Need to Differentiate between Bioadsorption and Bioaccumulation. New Biotechnol. 2012, 30 (1), 3–8. https://doi.org/10.1016/j.nbt.2012.05.020.

(34) Jeffrey, S. W.; Humphrey, G. F. New Spectrophotometric Equations for Determining Chlorophylls a, b, C1 and C2 in Higher Plants, Algae and Natural Phytoplankton. Biochem. Physiol. Pflanz. 1975, 167 (2), 191–194. https://doi.org/10.1016/S0015-3796(17)30778-3.

(35) Le Moigne, D.; Demay, J.; Reinhardt, A.; Bernard, C.; Kim Tiam, S.; Marie, B. Dynamics of the Metabolome of Aliinostoc Sp. PMC 882.14 in Response to Light and Temperature Variations. Metabolites 2021, 11 (11), 745. https://doi.org/10.3390/metabo11110745.

(36) Leguay, S.; Lavoie, I.; Levy, J. L.; Fortin, C. Using Biofilms for Monitoring Metal Contamination in Lotic Ecosystems: The Protective Effects of Hardness and PH on Metal Bioaccumulation: Monitoring Metal Contamination Using Stream Biofilms. Environ. Toxicol. Chem. 2016, 35 (6), 1489–1501. https://doi.org/10.1002/etc.3292.

(37) Niederer, C.; Behra, R.; Harder, A.; Schwarzenbach, R. P.; Escher, B. I. Mechanistic Approaches for Evaluating the Tocicity of Reactive Organochlorines and Epoxides in Green Algae. Environ. Toxicol. Chem. 2004, 23 (3), 697. https://doi.org/10.1897/03-83.

(38) Yu, X.; Jin, X.; Liu, H.; Yu, Y.; Tang, J.; Zhou, R.; Yin, A.; Sun, J.; Zhu, L. Enhanced Degradation of Atrazine through UV/Bisulfite: Mechanism, Reaction Pathways and Toxicological Analysis. Sci. Total Environ. 2023, 856, 159157. https://doi.org/10.1016/j.scitotenv.2022.159157.

(39) Morelli, E.; Scarano, G. Synthesis and Stability of Phytochelatins Induced by Cadmium and Lead in the Marine Diatom Phaeodactylum Tricornutum. Mar. Environ. Res. 2001, 52 (4), 383–395. https://doi.org/10.1016/S0141-1136(01)00093-9.

(40) Morelli, E.; Scarano, G. Copper-Induced Changes of Non-Protein Thiols and Antioxidant Enzymes in the Marine Microalga Phaeodactylum Tricornutum. Plant Sci. 2004, 167 (2), 289–296. https://doi.org/10.1016/j.plantsci.2004.04.001.

(41) Liu, W.; Majumdar, S.; Li, W.; Keller, A. A.; Slaveykova, V. I. Metabolomics for Early Detection of Stress in Freshwater Alga Poterioochromonas Malhamensis Exposed to Silver Nanoparticles. Sci. Rep. 2020, 10 (1), 20563. https://doi.org/10.1038/s41598-020-77521-0.

(42) Slaveykova, V. I.; Majumdar, S.; Regier, N.; Li, W.; Keller, A. A. Metabolomic Responses of Green Alga *Chlamydomonas Reinhardtii* Exposed to Sublethal Concentrations of Inorganic and Methylmercury. Environ. Sci. Technol. 2021, 55 (6), 3876–3887. https://doi.org/10.1021/acs.est.0c08416.

(43) Nishikawa, K.; Yamakoshi, Y.; Uemura, I.; Tominaga, N. Ultrastructural Changes in Chlamydomonas Acidophila (Chlorophyta) Induced by Heavy Metals and Polyphosphate Metabolism. FEMS Microbiol. Ecol. 2003, 44 (2), 253–259. https://doi.org/10.1016/S0168-6496(03)00049-7.

(44) Zhang, S.; He, Y.; Sen, B.; Wang, G. Reactive Oxygen Species and Their Applications toward Enhanced Lipid Accumulation in Oleaginous Microorganisms. Bioresour. Technol. 2020, 307, 123234. https://doi.org/10.1016/j.biortech.2020.123234.

(45) Popko, J. Lipid Composition of Chlamydomonas Reinhardtii. In *Encyclopedia of Lipidomics*; Wenk, M. R., Ed.; Springer Netherlands: Dordrecht, 2016; pp 1–6. https://doi.org/10.1007/978-94-007-7864-1_126-1.

(46) Fadhlaoui, M.; Laderriere, V.; Lavoie, I.; Fortin, C. Influence of Temperature and Nickel on Algal Biofilm Fatty Acid Composition. Environ. Toxicol. Chem. 2020, 39 (8), 1566–1577. https://doi.org/10.1002/etc.4741.

(47) Yang, P.; Pan, J.; Wang, H.; Xiaohan, X.; Zeling, X.; Chen, X.; Yang, Y.; Sun, H.; Li, M.; Habimana, O. Structural, Metagenomic and Metabolic Shifts in Multispecies Freshwater Biofilm Models Exposed to Silver Nanoparticles. J. Environ. Chem. Eng. 2023, 11 (1), 109162. https://doi.org/10.1016/j.jece.2022.109162.

(48) Favre, L.; Ortalo-Magné, A.; Kerloch, L.; Pichereaux, C.; Misson, B.; Briand, J.-F.; Garnier, C.; Culioli, G. Metabolomic and Proteomic Changes Induced by Growth Inhibitory Concentrations of Copper in the Biofilm-Forming Marine Bacterium *Pseudoalteromonas Lipolytica*. Metallomics 2019, 11 (11), 1887–1899. https://doi.org/10.1039/C9MT00184K.

(49) Massieux, B.; Boivin, M. E. Y.; van den Ende, F. P.; Langenskiöld, J.; Marvan, P.; Barranguet, C.; Admiraal, W.; Laanbroek, H. J.; Zwart, G. Analysis of Structural and Physiological Profiles To Assess the Effects of Cu on Biofilm Microbial Communities. Appl. Environ. Microbiol. 2004, 70 (8), 4512–4521. https://doi.org/10.1128/AEM.70.8.4512-4521.2004.

(50) Corcoll, N.; Bonet, B.; Leira, M.; Guasch, H. Chl-a Fluorescence Parameters as Biomarkers of Metal Toxicity in Fluvial Biofilms: An Experimental Study. Hydrobiologia 2011, 673 (1), 119–136. https://doi.org/10.1007/s10750-011-0763-8.

(51) Plekhanov, S. E.; Chemeris, Y. K. Early Toxic Effects of Zinc, Cobalt, and Cadmium on Photosynthetic Activity of the Green Alga Chlorella Pyrenoidosa Chick S-39. Biol. Bull. Russ. Acad. Sci. 2003, 30 (5), 6. https://doi.org/10.1023/A:1025806921291.

(52) El-Din, S. M. M. Effects of Heavy Metals (Copper, Cobalt and Lead) on the Growth and Photosynthetic Pigments of the Green Alga Chlorella Pyrenoidosa H. Chick. Catrina Int. J. Environ. Sci. 2016, 15 (1), 1–10.

(53) Jeffrey, S. W. Profiles of Photosynthetic Pigments in the Ocean Using Thin-Layer Chromatography. Mar. Biol. 1974, 26 (2), 101–110. https://doi.org/10.1007/BF00388879.

(54) Motten, A. F. Diversity of Photosynthetic Pigments. In Tested studies for laboratory teaching; Association fo rBiology Laboratory Education (ABLE): Las Vegas, 2004; pp 159–177.

(55) Aditi, N.; Jaishini, C.; Raisa, F.; Aamna, K.; Rupali, B. Photosynthetic Pigments, Lipids and Phenolic Compounds of Three Green Algae Isolated from Freshwater Ecosystem. J. Algal Biomass Util. 2020, 11, 68–83.

(56) Alberte, R. S.; Friedman, A. L.; Gustafson, D. L.; Rudnick, M. S.; Lyman, H. Light-Harvesting Systems of Brown Algae and Diatoms. Isolation and Characterization of Chlorophyll Ac and Chlorophyll Afucoxanthin Pigment-Protein Complexes. Biochim. Biophys. Acta BBA - Bioenerg. 1981, 635 (2), 304–316. https://doi.org/10.1016/0005-2728(81)90029-3.

(57) Harker, M.; Berkaloff, C.; Lemoine, Y.; Britton, G.; Young, A.; Duval, J.-C.; Rmiki, N.- E.; Rousseau, B. Effects of High Light and Desiccation on the Operation of the Xanthophyll Cycle in Two Marine Brown Algae. Eur. J. Phycol. 1999, 34 (1), 35–42. https://doi.org/10.1080/09670269910001736062.

(58) Crump, K. S. A New Method for Determining Allowable Daily Intakes. Toxicol. Sci. 1984, 4 (5), 854–871. https://doi.org/10.1016/0272-0590(84)90107-6.

(59) Gaylor, D.; Ryan, L.; Krewski, D.; Zhu, Y. Procedures for Calculating Benchmark Doses for Health Risk Assessment. Regul. Toxicol. Pharmacol. 1998, 28 (2), 150–164. https://doi.org/10.1006/rtph.1998.1247.

(60) Bokkers, B. G. H.; Slob, W. A Comparison of Ratio Distributions Based on the NOAEL and the Benchmark Approach for Subchronic-to-Chronic Extrapolation. Toxicol. Sci. 2005, 85 (2), 1033–1040. https://doi.org/10.1093/toxsci/kfi144.

(61) Bokkers, B. G. H.; Slob, W. Deriving a Data-Based Interspecies Assessment Factor Using the NOAEL and the Benchmark Dose Approach. Crit. Rev. Toxicol. 2007, 37 (5), 355–373. https://doi.org/10.1080/10408440701249224.

(62) Bonvallot, N.; Bodin, L.; Duboudin, C.; Bard, D. Benchmark dose: définitions, intérêt et usages en évaluation des risques sanitaires. *Environ. Risques Santé* 2009, 8 (6).

(63) Mayfield, D. B.; Skall, D. G. Benchmark Dose Analysis Framework for Developing Wildlife Toxicity Reference Values: BMD Analysis Framework for Wildlife TRVs. Environ. Toxicol. Chem. 2018, 37 (5), 1496–1508. https://doi.org/10.1002/etc.4082.

(64) Stubblefield, W. A.; Van Genderen, E.; Cardwell, A. S.; Heijerick, D. G.; Janssen, C. R.; De Schamphelaere, K. A. C. Acute and Chronic Toxicity of Cobalt to Freshwater Organisms: Using a Species Sensitivity Distribution Approach to Establish International Water Quality Standards. Environ. Toxicol. Chem. 2020, 39 (4), 799–811. https://doi.org/10.1002/etc.4662.

(65) Cobbett, C.; Goldsbrough, P. Phytochelatins and Metallothioneins: Roles in Heavy Metal Detoxification and Homeostasis. Annu. Rev. Plant Biol. 2002, 53 (1), 159–182. https://doi.org/10.1146/annurev.arplant.53.100301.135154.

(66) Hall, J. L. Cellular Mechanisms for Heavy Metal Detoxification and Tolerance. J. Exp. Bot. 2002, 53 (366), 1–11. https://doi.org/10.1093/jexbot/53.366.1.

(67) Foyer, C. H.; Noctor, G. Redox Homeostasis and Antioxidant Signaling: A Metabolic Interface between Stress Perception and Physiological Responses. Plant Cell 2005, 17 (7), 1866–1875. https://doi.org/10.1105/tpc.105.033589.

(68) Torres, M. A.; Barros, M. P.; Campos, S. C. G.; Pinto, E.; Rajamani, S.; Sayre, R. T.; Colepicolo, P. Biochemical Biomarkers in Algae and Marine Pollution: A Review. Ecotoxicol. Environ. Saf. 2008, 71 (1), 1–15. https://doi.org/10.1016/j.ecoenv.2008.05.009.

(69) Chen, M.; Tang, H.; Ma, H.; Holland, T. C.; Ng, K. Y. S.; Salley, S. O. Effect of Nutrients on Growth and Lipid Accumulation in the Green Algae Dunaliella Tertiolecta. Bioresour. Technol. 2011, 102 (2), 1649–1655. https://doi.org/10.1016/j.biortech.2010.09.062.

(70) Bártová, K.; Hilscherová, K.; Babica, P.; Maršálek, B.; Bláha, L. Effects of Microcystin and Complex Cyanobacterial Samples on the Growth and Oxidative Stress Parameters in Green Alga Pseudokirchneriella Subcapitata and Comparison with the Model Oxidative Stressor-Herbicide Paraquat. Environ. Toxicol. 2011, 26 (6), 641–648. https://doi.org/10.1002/tox.20601.

(71) Hong, Y.; Tan, Y.; Meng, Y.; Yang, H.; Zhang, Y.; Warren, A.; Li, J.; Lin, X. Evaluation of Biomarkers for Ecotoxicity Assessment by Dose-Response Dynamic Models: Effects of Nitrofurazone on Antioxidant Enzymes in the Model Ciliated Protozoan Euplotes Vannus. Ecotoxicol. Environ. Saf. 2017, 144, 552–559. https://doi.org/10.1016/j.ecoenv.2017.06.069.

(72) Abassi, S.; Wang, H.; Ponmani, T.; Ki, J. Small Heat Shock Protein Genes of the Green Algae *Closterium Ehrenbergii* : Cloning and Differential Expression under Heat and Heavy Metal Stresses. Environ. Toxicol. 2019, 34 (9), 1013–1024. https://doi.org/10.1002/tox.22772.

(73) Nie, X.; Wang, X.; Chen, J.; Zitko, V.; An, T. Response of the Freshwater Alga Chlorella Vulgaris to Trichloroisocyanuric Acid and Ciprofloxacin. Environ. Toxicol. Chem. 2008, 27 (1), 168. https://doi.org/10.1897/07-028.1.

(74) Melegari, S. P.; Perreault, F.; Costa, R. H. R.; Popovic, R.; Matias, W. G. Evaluation of Toxicity and Oxidative Stress Induced by Copper Oxide Nanoparticles in the Green Alga Chlamydomonas Reinhardtii. Aquat. Toxicol. 2013, 142–143, 431–440. https://doi.org/10.1016/j.aquatox.2013.09.015.

(75) Xiao, A.; Wang, C.; Chen, J.; Guo, R.; Yan, Z.; Chen, J. Carbon and Metal Quantum Dots Toxicity on the Microalgae Chlorella Pyrenoidosa. Ecotoxicol. Environ. Saf. 2016, 133, 211–217. https://doi.org/10.1016/j.ecoenv.2016.07.026.

(76) Fang, B.; Shi, J.; Qin, L.; Feng, M.; Cheng, D.; Wang, T.; Zhang, X. Toxicity Evaluation of 4,4′-Di-CDPS and 4,4′-Di-CDE on Green Algae Scenedesmus Obliquus: Growth Inhibition, Change in Pigment Content, and Oxidative Stress. Environ. Sci. Pollut. Res. 2018, 25 (16), 15630– 15640. https://doi.org/10.1007/s11356-018-1749-0.

(77) Çelekli, A.; Kapı, M.; Bozkurt, H. Effect of Cadmium on Biomass, Pigmentation, Malondialdehyde, and Proline of Scenedesmus Quadricauda Var. Longispina. Bull. Environ. Contam. Toxicol. 2013, 91 (5), 571–576. https://doi.org/10.1007/s00128-013-1100-x.

(78) Rai, U. N.; Singh, N. K.; Upadhyay, A. K.; Verma, S. Chromate Tolerance and Accumulation in Chlorella Vulgaris L.: Role of Antioxidant Enzymes and Biochemical Changes in Detoxification of Metals. Bioresour. Technol. 2013, 136, 604–609. https://doi.org/10.1016/j.biortech.2013.03.043.

(79) Fawzy, M. A.; Hifney, A. F.; Adam, M. S.; Al-Badaani, A. A. Biosorption of Cobalt and Its Effect on Growth and Metabolites of Synechocystis Pevalekii and Scenedesmus Bernardii: Isothermal Analysis. *Environ*. Technol. Innov. 2020, 19, 100953. https://doi.org/10.1016/j.eti.2020.100953.

