## Supporting Information for "Meta-metabolomic Responses of River Biofilms to Cobalt Exposure and Use of Dose-response Model Trends as an Indicator of Effects"

For

*Corresponding author:

**CONTENTS**

**Texts**

**Text S1.** Biofilm colonization and exposure to Co in microcosms.

**Text S2.** Water analysis and Co speciation.

**Text S3.** Total and intracellular Co concentration.

**Text S4.** Chlorophyll analysis.

**Text S5.** Meta-metabolomic analysis.

**Text S6.** Metal bioaccumulation.

**Text S7.** Effect of Co on annotated metabolites.

**Tables (see .xlsx files)**

**Table S1.** Physico-chemical parameters of the *Gave de Pau* water.

**Table S2.** Physico-chemical parameters of the microcosms.

**Table S3.** Total and intracellular concentrations of metals (mean and sd).

**Table S4.** Raw metabolomic data set.

**Table S5.** PLS-DA cross-validation details.

**Table S6.** ECDF parameters.

**Figures**

**Figure S1.** Benchmark-dose calculation illustration.

**Figure S2.** Cobalt bioaccumulation of biofilms. Total and intracellular Co concentrations ($log_{10}$ $mol \cdot g^{-1}_{DW}$) as function of $Co^{2+}$ concentration (log10 M) in exposure medium at different exposure time.

**Figure S3.** Meta-metabolomic response of biofilms according to time and concentrations of exposure to Co. (A) Individual score plot generated from PLS-DA analysis performed with 159 annotated variables to components 1-2 with all conditions grouped together, (B) after one day of exposure, (C) three and (D) seven. Quality descriptors of the statistical significance and predictive ability of the discriminant model according to (E) cross-validations test and (F) corresponding permutations. (G) Four examples of representative box-plot. (H) Heatmap with hierarchical classification representation of concentration class averages (Ward clustering according to Euclidian distances) performed from relative intensities of the dysregulated analytes with a VIP score > 1 for the component 1.

**Figure S4.** Four examples of representative box plots of metabolite fluctuations

**Figure S5.** t-distributed Stochastic Neighbor Embedding of annotated metabolites.

#### Texts

**Text S1.** Biofilm colonization and exposure to Co in microcosms.

The TotalEnergies facility in Pau (PERL, France) consists of open mesocosms (0.5 m x 0.5 m x 40 m) and a *so-called* nursery situated upstream the mesocosms in which the *Gave de Pau* naturally flows. The bottom of that nursery is made of rocks and is naturally colonized by river organisms (biofilms, plants, mollusks and small fishes). Biofilm were colonized in that nursery.

One day before the start of the exposure (D-1), 15 acid-rinsed plastic microcosms (29 cm x 39 cm x 26 cm) were filled with 15 L of the *Gave de Pau* water. A total of three microcosms for each exposure conditions (background concentration (control; 2 x $10^{-9}$ M), 1 x $10^{-7}$, 1 x $10^{-6}$, 5 x $10^{-6}$ and 1 x $10^{-5}$ M Co – precisely determined) were placed directly in one mesocosm in order to maintain the water temperature at 13.5 ± 1°C (Table S1). Cobalt (Cobalt Standard for ICP, 1000 mg·$L^{-1}$, Supelco, Germany) was added in each microcosm to let equilibrating for 24 h. After that addition, the pHs of the exposure media were adjusted with NaOH 1 M (99.99%, Sigma-Aldrich, Germany) or $HNO_3$ 70% (≥99.999% trace metal basis, Sigma-Aldrich, Germany) in order to maintain them within similar values than those of the mesocosm stream water (7.9 ± 0.2, n = 6). One hour after the first addition (D-1), water was collected for further analysis. During the exposure experiments (seven days in total), additional Co was added to the exposure media at D2, D3 and D6 (except for 5 x $10^{-6}$ and 1 x $10^{-5}$ M Co) to maintain Co concentrations. Each time, pHs were adjusted when necessary. Water (D-1, D0, D1, D3, and D7) were collected prior Co spiking and pH adjustment.

The first day of exposure experiment (D0), seven biofilm colonized slides were placed in each microcosm (105 in total). Additionally, eight slides were directly taken from the nursery as a D0 control; three to study Co accumulation (one side for total concentration and the second side for intracellular concentration), three for meta-metabolomics, two for chlorophyll content. Each side of the slide was scraped with a microscope glass slide, placed separately in 2 mL Eppendorfs® tubes and stored in the dark at -20°C.

**Text S2.** Water analysis and Co speciation.

Water was filtered with a syringe (Solo Injekt®, Braun, Germany) equipped with a 0.45 µm polysulfone filter (Minisart®, Supelco, Germany) and collected in 15 mL-polypropylene (PP) Falcon™ metal free, Labcon, CA, USA) tubes. Metal and cation samples were acidified with 200 µL $HNO_3$ 70% and placed at 4°C in the dark. Anion samples were not acidified but stored under similar conditions. Three times 125 mL of water were also filtered at D-1 and stored in amber bottles previously burnt at 500°C for 4 h for DOC analysis. Those DOC samples were acidified with 200 µL of HCl 2 M and stored at 4°C in the dark. Water temperature and pH were measured using a multi-parameters field probe (Hach, IA, USA). Metal concentrations were analyzed by inductively Coupled Plasma Mass Spectrometry (ICP-MS model 7500, Agilent, CA, USA). Anion and cation concentrations were analysed by ion chromatography (Dionex Aquion, Thermo-Fisher, MA, USA). The accuracy of both techniques was verified using SRL-6 (River water, NRC Canada) as certified reference material. The percentage of Co recovery was 99.3 ± 0.1 % (n=3). The DOC concentrations were analyzed using a ShimatzuTOC-L, analyzer (Japan). Co speciation was modelled using the Windermere Humic Aqueous Model (WHAM) model VII with its default binding constants. A ratio of 2 was used between DOC and the dissolved organic matter (DOM) concentration, whereas the DOM composition was estimated to be composed of 65% of fulvic acid, the rest being inert.[1,2]

**Text S3.** Total and intracellular Co concentration.

To determine the total Co concentration in biofilms, biofilm samples were lyophilized during 24 h then weighted (~ 60 mg) in quartz vials (Pyrex ®, France)) to which 3.3 mL $HNO_3$ 70% and 1.7 mL $H_2O_2$ 30% (w/w, for ultratrace analysis, Supelco, Germany) were added. After these additions, the vials were placed in an UltraWAVE™ oven (Milestone, Italy). Temperature and pressure were raised to 220°C and 110 bar, respectively, for 25 minutes. The digestates were decanted in Falcon™ metal free tubes, weighted and diluted in Milli-Q water. A centrifugation (5000 rpm; 5 min) was then performed to remove any remaining particles. The supernatant was further diluted with Milli-Q water to obtain a final concentration of 2% $HNO_3$. To distinguish between total and intracellular Co content, biofilms were previously additionally rinsed with 10 mM ethylenediaminetetraacetic acid (EDTA, under disodium salt dihydrate form, Sigma-Aldrich, Germany) for 10 min in order to remove adsorbed metals on the biofilm surface. After that rinsing step, the biofilm samples for intracellular concentration were freeze-dried and processed similarly as samples for the total concentration analysis. Metal concentrations were measured by ICP-MS (ICP-MS model 7500, Agilent, CA, USA). The digestion accuracy was verified using BCR-414 (plankton, JRC, Brussels) as certified reference material. The percentage of Co recovery was 91 ± 8 % (n=9).

**Text S4.** Chlorophyll analysis.

For the determination of the chlorophyll content, 10 mL of 90% acetone (Uvasol®, Sigma-Aldrich, Germany) were added to biofilm samples (50 $cm^2$ of sample), sonicated at 30% for 30 min and left in the dark for 12 h. The samples were then centrifuged (6000 g, 10 min), and 1 mL of supernatant was analyzed by spectrophotometry (Lambda 750, PerkinElmer, MA, USA). Different wavelengths between 400 and 750 nm (with a step of 1 nm) were used to quantify the chlorophyll *a*, chlorophyll *b* and chlorophylls $c_1 + c_2$ content following the Jeffrey and Humphrey protocol.[3]

**Text S5.** Meta-metabolomic analysis.

The samples were firstly analyzed in simple MS positive mode, without quadrupole fragmentation to obtain a measured peak intensity for each ion signal (measured in number of detected counts over the time range in the different samples). MetaboScape 4.0 software (Bruker, Bremen, Germany) was then used to recalibrate the obtained MS data. Only ions that had an intensity greater than 5000 counts in at least 10% of the set of samples were selected. Thereafter autoMS/MS analysis was performed in positive mode for qualitative investigation of metabolites. Ions with the highest intensities in single MS were selected and fragmentated by collision ion dissociation (CID). The obtained MS/MS data file (.mgf file) was used to annotate the metabolites and to generate a molecular network for comparison of fragmentation profiles using the MetGem 1.3.6. and the GNPS algorithm. Subsequently different spectral databases were used, i.e., GNPS library, NIH Clinical collections, and EMBL metabolomics to annotate as much metabolites as possible.

**Text S6.** Metal bioaccumulation.

Biofilm naturally contained 1.3 ± 0.1 x 10-7 mol·$g_{DW}^{-1}$ total accumulated Co (Table S3). These concentrations significantly increased upon exposure to increasing Co concentrations (Figure 1A and Table S3) at each sampling time (D1, D3 and D7). For example, at D7, the total Co bioaccumulated concentration increased from 7.4 ± 1.1 x $10^{-8}$ mol·$g_{DW}^{-1}$ (control), to 1.2 ± 0.2 x $10^{-5}$ mol·$g_{DW}^{-1}$ and 3.7 ± 1.1 x $10^{-5}$ mol·$g_{DW}^{-1}$ upon exposure to 1 x $10^{-6}$ M and 1 x $10^{-5}$ M Co, respectively. Total Co bioaccumulation was also found to increase with exposure time (Kruskal-Wallis test, p-value: < 0.05) (FigureS2 and Table S3). As such, the total Co accumulation was greater at D7 than at D1 and D3 (*post-hoc* Dunn test, p-value D1 – D3 > 0.05; p-value D1 -D7 < 0.05; p-value D3 – D7 < 0.05). In contrast, a progressive decrease in the concentrations of bioaccumulated Co over time in the control condition was observed, from 1.4 x $10^{-7}$ M to 5.4 x $10^{-8}$ M between D1 and D7. (Figure S1). The accumulation of total Co as a function of $Co^{2+}$ concentration ($log_{10}$) was linear at each exposure time with an increase in slopes from 0.5 at D1, to 0.55 at D3 and 0.77 at D7 (Figure S1). The $R^2_{adj}$ also increased from 0.7 to 0.95 between D1 and D7.

Similarly to total Co bioaccumulation, the intracellular Co bioaccumulation was significantly correlated with total dissolved Co and $Co^{2+}$ concentrations in the exposure media (Figure 1A, Table S3). Moreover, for similar $Co^{2+}$ concentrations in the media, intracellular Co concentration was significantly different between each day (*post-hoc* Dunn test, p-values < 0.05). At D7, that concentration was measured to be greater than the one at D3, which itself was greater than the one at D1 (*post-hoc* Dunn tests, D1 – D3 < 0.05; D1 – D7 < 0.05; D3 – D7 < 0.05). The accumulation of intracellular Co as a function of $Co^{2+}$ concentration ($log_{10}$) was also linear at each exposure time with an increase in slopes from 0.43 at D1, to 0.58 at D3 and 0.74 at D7 (Figure S1). The $R^2_{adj}$ also increased from 0.91 to 0.96 between D1 and D7.

**Text S7.** Effect of Co on annotated metabolites.

As for non-annotated metabolites, PLS-DA was constructed to compare treatments. That PLS-DA model showed significant $R^2$ *cumulative*, $Q^2$ *cumulative* and permutation score (Fig. S2E, S2F). The meta-metabolomic response of the control biofilms was different at each studied time (PERMANOVA, p-value < 0.05). At D1, Co had no significant impact on the annotated meta-metabolomic signature of biofilms (Fig. S2B). At D3, differences in metabolic response were observed as a function of the Co concentration in the media. Biofilms exposed to natural concentrations ($2 \times 10^{-9}$ M), $1 \times 10^{-7}$ and $1 \times 10^{-6}$ M had the same meta-metabolomic profile, which was significantly different from those of biofilms exposed to $5 \times 10^{-6}$ and $1 \times 10^{-5}$ M Co (PERMANOVA, p-value < 0.05). Biofilms exposed to these two highest concentrations had their meta-metabolomic responses different from each other (PERMANOVA, p-value < 0.01). At D7, only biofilms exposed to the control and $1 \times 10^{-7}$ M Co had a similar metabolic profile (PERMANOVA, p-values < 0.05). For each exposure condition, the annotated meta-metabolomic signature of the biofilms was different between D1 and D7 (PERMANOVA, p-value < 0.05).

### Tables

**Table S1.** Physico-chemical parameters of the *Gave de Pau* water.

| Co exposure concentration (M) | Replicate | Date | Hour | Day | Temperature (°C) | pH | Total Co (M) | $Co^{2+}$ (M) | Ca (M) | K (M) | Mg (M) | Na (M) | Sulfates ($mg.L^{-1}$) | Nitrates ($mg.L^{-1}$) | Chlorides ($mg.L^{-1}$) | DOC ($mg.L^{-1}$) |
|---|---|---|---|---|---|---|---|---|---|---|---|---|---|---|---|---|
| Gave de Pau | 6 | 03/05/2021 | 10:00 a.m. | D-1 | 10.50 ± 0.00 | 7.90 ± 0.00 | 1.71E-9 ± 1.88E-10 | 1.09E-9 ± 1.24E-10 | 4.07E-5 ± 5.27E-7 | 1.02E-6 ± 1.09E-8 | 2.87E-6 ± 4.52E-8 | 2.07E-6 ± 2.32E-8 | 12.10 ± 0.11 | 3.33 ± 0.34 | 5.4 ± 0.03 | 1.22 ± 0.28 |

**Table S2.** Physico-chemical parameters of the microcosms.

| Co exposure concentration (M) | Replicate | Date | Hour | Day | Temperature (°C) | pH | Total Co (M) | $Co^{2+}$ (M) |
|---|---|---|---|---|---|---|---|---|
| 0 | 3 | 05/05/2021 | 10:00 a.m. | D1 | 14.27 ± 0.06 | 7.97 ± 0.04 | 2.62E-9 ± 3.97E-10 | 1.46E-9 ± 1.46E-10 |
| 1 x $10^{-7}$ | 3 | 05/05/2021 | 10:05 a.m. | D1 | 14.43 ± 0.12 | 7.92 ± 0.09 | 6.88E-8 ± 2.78E-9 | 4.13E-8 ± 6.26E-9 |
| 1 x $10^{-6}$ | 3 | 05/05/2021 | 10:10 a.m. | D1 | 14.43 ± 0.06 | 7.93 ± 0.09 | 7.33E-7 ± 4.09E-8 | 4.38E-7 ± 7.72E-8 |
| 5 x $10^{-6}$ | 3 | 05/05/2021 | 10:15 a.m. | D1 | 14.37 ± 0.12 | 7.77 ± 0.02 | 4.41E-6 ± 5.88E-8 | 3.16E-6 ± 3.61E-8 |
| 1 x $10^{-5}$ | 3 | 05/05/2021 | 10:20 a.m. | D1 | 14.40 ± 0.10 | 7.90 ± 0.06 | 8.09E-6 ± 2.28E-7 | 5.06E-6 ± 5.24E-7 |
| 0 | 3 | 07/05/2021 | 10:00 a.m. | D3 | 13.97 ± 0.06 | 8.07 ± 0.14 | 1.45E-9 ± 5.23E-11 | 6.80E-10 ± 1.94E-10 |
| 1 x $10^{-7}$ | 3 | 07/05/2021 | 10:05 a.m. | D3 | 14.00 ± 0.00 | 8.12 ± 0.04 | 1.25E-7 ± 4.81E-9 | 5.29E-8 ± 5.43E-9 |
| 1 x $10^{-6}$ | 3 | 07/05/2021 | 10:10 a.m. | D3 | 13.93 ± 0.06 | 8.31 ± 0.15 | 1.38E-6 ± 1.30E-7 | 3.71E-7 ± 1.50E-7 |
| 5 x $10^{-6}$ | 3 | 07/05/2021 | 10:15 a.m. | D3 | 13.90 ± 0.00 | 7.94 ± 0.04 | 8.60E-6 ± 1.28E-7 | 5.07E-6 ± 2.21E-7 |
| 1 x $10^{-5}$ | 3 | 07/05/2021 | 10:20 a.m. | D3 | 14.00 ± 0.00 | 7.98 ± 0.07 | 1.59E-5 ± 2.52E-7 | 8.82E-6 ± 1.00E-6 |
| 0 | 3 | 11/05/2021 | 10:00 a.m. | D7 | 11.90 ± 0.00 | 8.09 ± 0.12 | 5.25E-9 ± 9.16E-10 | 2.43E-9 ± 7.75E-10 |
| 1 x $10^{-7}$ | 3 | 11/05/2021 | 10:05 a.m. | D7 | 12.00 ± 0.00 | 8.04 ± 0.23 | 2.37E-7 ± 5.88E-8 | 1.26E-7 ± 8.17E-8 |
| 1 x $10^{-6}$ | 3 | 11/05/2021 | 10:10 a.m. | D7 | 12.07 ± 0.06 | 8.03 ± 0.09 | 2.59E-6 ± 2.01E-7 | 1.36E-6 ± 3.11E-7 |
| 5 x $10^{-6}$ | 3 | 11/05/2021 | 10:15 a.m. | D7 | 12.07 ± 0.06 | 8.28 ± 0.09 | 1.28E-5 ± 4.34E-7 | 3.76E-6 ± 8.00E-7 |
| 1 x $10^{-5}$ | 3 | 11/05/2021 | 10:20 a.m. | D7 | 13.50 ± 0.00 | 8.14 ± 0.07 | 2.41E-5 ± 3.78E-7 | 1.00E-5 ± 1.52E-6 |

**Table S3.** Total and intracellular concentrations of metals (mean and sd).

| Bioaccumulated concentration | Co exposure concentration (M) | Replicate | Date | Hour | Day | Co $(mol \cdot g_{DW}^{-1})$ | Li $(mol \cdot g_{DW}^{-1})$ | Ni $(mol \cdot g_{DW}^{-1})$ | Cu $(mol \cdot g_{DW}^{-1})$ | Zn $(mol \cdot g_{DW}^{-1})$ | As $(mol \cdot g_{DW}^{-1})$ | Pb $(mol \cdot g_{DW}^{-1})$ |
|---|---|---|---|---|---|---|---|---|---|---|---|---|
| Intracellular | control | 3 | 03/02/2021 | 10:05 a.m. | D-1 | 1.23E-7 ± 8.95E-9 | 6.01E-6 ± 5.06E-7 | 2.21E-6 ± 2.30E-7 | 3.78E-7 ± 9.60E-8 | 4.32E-6 ± 8.18E-7 | 2.11E-7 ± 2.88E-9 | NA ± NA |
| Intracellular | 0 | 3 | 05/05/2021 | 10:25 a.m. | D1 | 1.41E-7 ± 9.92E-9 | 5.46E-6 ± 1.69E-7 | 2.04E-6 ± 1.51E-7 | 2.69E-7 ± 4.82E-8 | 3.21E-6 ± 2.38E-7 | 1.63E-7 ± 1.59E-8 | 1.52E-7 ± 6.95E-9 |
| Intracellular | $1 \times 10^{-7}$ | 3 | 05/05/2021 | 10:30 a.m. | D1 | 2.55E-7 ± 4.05E-8 | 6.47E-6 ± 4.94E-7 | 2.07E-6 ± 4.43E-7 | 2.61E-7 ± 5.78E-8 | 3.16E-6 ± 6.14E-7 | 1.95E-7 ± 2.84E-8 | 1.59E-7 ± 3.09E-8 |
| Intracellular | $1 \times 10^{-6}$ | 3 | 05/05/2021 | 10:35 a.m. | D1 | 1.44E-6 ± 7.22E-7 | 5.71E-6 ± 5.94E-7 | 1.92E-6 ± 3.59E-7 | 2.94E-7 ± 4.56E-8 | 4.20E-6 ± 4.15E-7 | 2.23E-7 ± 1.44E-8 | 2.13E-7 ± 1.49E-8 |
| Intracellular | $5 \times 10^{-6}$ | 3 | 05/05/2021 | 10:40 a.m. | D1 | 2.40E-6 ± 4.36E-7 | 5.49E-6 ± 3.97E-7 | 2.05E-6 ± 1.07E-7 | 2.49E-7 ± 1.85E-8 | 2.90E-6 ± 3.65E-7 | 1.67E-7 ± 1.21E-8 | 1.69E-7 ± 2.51E-8 |
| Intracellular | $1 \times 10^{-5}$ | 3 | 05/05/2021 | 10:45 a.m. | D1 | 5.07E-6 ± 1.41E-6 | 5.66E-6 ± 3.79E-7 | 1.98E-6 ± 1.94E-7 | 2.55E-7 ± 4.89E-8 | 3.10E-6 ± 5.52E-7 | 1.91E-7 ± 2.11E-8 | 1.54E-7 ± 1.52E-8 |
| Intracellular | 0 | 3 | 07/05/2021 | 10:25 a.m. | D3 | 7.46E-8 ± 3.91E-8 | 3.69E-6 ± 1.12E-6 | 1.18E-6 ± 4.81E-7 | 1.54E-7 ± 7.25E-8 | 2.43E-6 ± 5.58E-7 | 1.54E-7 ± 2.28E-8 | 1.12E-7 ± 2.88E-8 |
| Intracellular | $1 \times 10^{-7}$ | 3 | 07/05/2021 | 10:30 a.m. | D3 | 5.69E-7 ± 2.86E-7 | 4.10E-6 ± 1.14E-6 | 1.49E-6 ± 4.75E-7 | 2.42E-7 ± 6.49E-8 | 4.81E-6 ± 3.76E-6 | 1.67E-7 ± 7.18E-9 | 1.34E-7 ± 1.28E-8 |
| Intracellular | $1 \times 10^{-6}$ | 3 | 07/05/2021 | 10:35 a.m. | D3 | 3.14E-6 ± 9.69E-7 | 4.95E-6 ± 2.85E-7 | 1.62E-6 ± 1.56E-7 | 1.99E-7 ± 3.12E-8 | 2.08E-6 ± 3.68E-7 | 1.53E-7 ± 1.82E-8 | 1.38E-7 ± 1.50E-8 |
| Intracellular | $5 \times 10^{-6}$ | 3 | 07/05/2021 | 10:40 a.m. | D3 | 8.69E-6 ± 2.11E-6 | 4.98E-6 ± 2.32E-7 | 1.29E-6 ± 7.99E-7 | 1.37E-7 ± 1.07E-7 | 2.54E-6 ± 2.63E-6 | 1.65E-7 ± 6.26E-9 | 1.35E-7 ± 5.60E-9 |
| Intracellular | $1 \times 10^{-5}$ | 3 | 07/05/2021 | 10:45 a.m. | D3 | 1.90E-5 ± 6.58E-6 | 2.93E-6 ± 1.92E-6 | 1.12E-6 ± 7.69E-7 | 1.18E-7 ± 1.01E-7 | 3.02E-6 ± 7.19E-7 | 1.27E-7 ± 3.22E-8 | 9.77E-8 ± 3.59E-8 |
| Intracellular | 0 | 3 | 11/05/2021 | 10:25 a.m. | D7 | 5.41E-8 ± 1.60E-8 | 2.65E-6 ± 5.27E-7 | 1.06E-6 ± 2.18E-7 | 9.72E-8 ± 2.03E-8 | 1.50E-6 ± 8.46E-7 | 1.22E-7 ± 2.40E-8 | 9.69E-8 ± 1.63E-8 |
| Intracellular | $1 \times 10^{-7}$ | 3 | 11/05/2021 | 10:30 a.m. | D7 | 1.41E-6 ± 3.93E-7 | 2.59E-6 ± 7.12E-7 | 8.09E-7 ± 4.04E-7 | 9.13E-8 ± 3.58E-8 | 1.71E-6 ± 2.80E-7 | 1.36E-7 ± 2.39E-8 | 8.99E-8 ± 1.44E-8 |
| Intracellular | $1 \times 10^{-6}$ | 3 | 11/05/2021 | 10:35 a.m. | D7 | 7.56E-6 ± 9.98E-7 | 3.13E-6 ± 9.45E-7 | 8.50E-7 ± 4.20E-7 | 1.18E-7 ± 4.52E-8 | 2.03E-6 ± 4.90E-7 | 1.21E-7 ± 1.84E-8 | 9.86E-8 ± 3.66E-8 |
| Intracellular | $5 \times 10^{-6}$ | 3 | 11/05/2021 | 10:40 a.m. | D7 | 1.91E-5 ± 3.69E-6 | 5.37E-6 ± 6.51E-7 | 1.65E-6 ± 5.55E-7 | 2.84E-7 ± NA | 2.49E-6 ± NA | 1.95E-7 ± 3.36E-8 | 1.13E-7 ± 6.47E-8 |
| Intracellular | $1 \times 10^{-5}$ | 3 | 11/05/2021 | 10:45 a.m. | D7 | 2.72E-5 ± 1.46E-5 | 4.67E-6 ± 2.73E-6 | 1.64E-6 ± 8.67E-7 | 1.56E-7 ± 6.38E-8 | 2.64E-6 ± 6.79E-7 | 1.71E-7 ± 3.50E-8 | 1.24E-7 ± 3.29E-8 |
| Total | control | 3 | 03/02/2021 | 10:05 a.m. | D-1 | 1.34E-7 ± 1.08E-8 | 5.74E-6 ± 3.54E-7 | 2.06E-6 ± 7.23E-8 | 2.90E-7 ± 6.69E-8 | 3.66E-6 ± 7.38E-7 | 2.03E-7 ± 4.09E-8 | NA ± NA |
| Total | 0 | 3 | 05/05/2021 | 10:25 a.m. | D1 | 1.45E-7 ± 1.79E-8 | 5.66E-6 ± 8.61E-7 | 1.96E-6 ± 4.23E-7 | 2.97E-7 ± 8.62E-8 | 4.67E-6 ± 1.60E-6 | 2.16E-7 ± 4.87E-8 | 2.17E-7 ± 4.76E-8 |
| Total | $1 \times 10^{-7}$ | 3 | 05/05/2021 | 10:30 a.m. | D1 | 3.69E-7 ± 1.25E-7 | 5.19E-6 ± 6.26E-7 | 1.74E-6 ± 1.64E-7 | 2.93E-7 ± 1.38E-8 | 5.47E-6 ± 1.32E-6 | 2.15E-7 ± 2.94E-8 | 2.35E-7 ± 1.99E-8 |
| Total | $1 \times 10^{-6}$ | 3 | 05/05/2021 | 10:35 a.m. | D1 | 2.02E-6 ± 1.63E-6 | 4.54E-6 ± 1.63E-6 | 1.55E-6 ± 5.71E-7 | 1.96E-7 ± 6.89E-8 | 4.32E-6 ± 9.40E-7 | 1.76E-7 ± 2.38E-8 | 1.92E-7 ± 3.46E-8 |
| Total | $5 \times 10^{-6}$ | 3 | 05/05/2021 | 10:40 a.m. | D1 | 7.11E-6 ± 2.45E-6 | 5.11E-6 ± 5.43E-7 | 1.76E-6 ± 1.56E-7 | 2.46E-7 ± 1.47E-8 | 4.98E-6 ± 8.24E-7 | 2.30E-7 ± 5.28E-8 | 2.35E-7 ± 2.75E-8 |
| Total | $1 \times 10^{-5}$ | 3 | 05/05/2021 | 10:45 a.m. | D1 | 1.26E-5 ± 1.03E-5 | 3.62E-6 ± 3.12E-6 | 1.23E-6 ± 1.04E-6 | 2.05E-7 ± 1.85E-7 | 2.34E-6 ± 1.99E-6 | 1.72E-7 ± 1.47E-7 | 1.80E-7 ± 1.61E-7 |
| Total | 0 | 3 | 07/05/2021 | 10:25 a.m. | D3 | 9.99E-8 ± 2.56E-8 | 3.90E-6 ± 1.12E-6 | 1.37E-6 ± 3.97E-7 | 1.81E-7 ± 3.37E-8 | 3.03E-6 ± 5.77E-7 | 1.63E-7 ± 1.28E-8 | 1.44E-7 ± 3.76E-8 |
| Total | $1 \times 10^{-7}$ | 3 | 07/05/2021 | 10:30 a.m. | D3 | 7.78E-7 ± 4.03E-7 | 4.13E-6 ± 2.36E-7 | 1.66E-6 ± 8.87E-8 | 2.40E-7 ± 7.76E-8 | 3.38E-6 ± 7.27E-7 | 1.95E-7 ± 4.02E-8 | 1.82E-7 ± 3.74E-8 |
| Total | $1 \times 10^{-6}$ | 3 | 07/05/2021 | 10:35 a.m. | D3 | 4.31E-6 ± 7.54E-7 | 4.29E-6 ± 5.05E-7 | 1.47E-6 ± 1.28E-7 | 1.38E-7 ± 3.65E-8 | 3.71E-6 ± 6.32E-7 | 1.69E-7 ± 1.89E-8 | 1.67E-7 ± 8.80E-9 |
| Total | $5 \times 10^{-6}$ | 3 | 07/05/2021 | 10:40 a.m. | D3 | 5.55E-5 ± 7.38E-5 | 3.57E-5 ± 5.26E-5 | 1.28E-5 ± 1.86E-5 | 1.62E-6 ± 2.42E-6 | 1.51E-5 ± 2.25E-5 | 1.11E-6 ± 1.60E-6 | 1.36E-6 ± 2.04E-6 |
| Total | $1 \times 10^{-5}$ | 3 | 07/05/2021 | 10:45 a.m. | D3 | 1.86E-5 ± 2.61E-5 | 4.43E-7 ± 3.04E-7 | 1.47E-7 ± 9.76E-8 | 4.59E-8 ± 5.65E-8 | 1.06E-6 ± 1.45E-6 | 6.10E-8 ± 7.54E-8 | 3.26E-8 ± 3.63E-8 |
| Total | 0 | 3 | 11/05/2021 | 10:25 a.m. | D7 | 7.42E-8 ± 1.06E-8 | 2.43E-6 ± 3.82E-7 | 9.18E-7 ± 1.05E-7 | 1.20E-7 ± 2.25E-8 | 2.11E-6 ± 2.54E-7 | 1.30E-7 ± 1.05E-8 | 9.46E-8 ± 9.16E-9 |
| Total | $1 \times 10^{-7}$ | 3 | 11/05/2021 | 10:30 a.m. | D7 | 1.73E-6 ± 3.60E-7 | 2.52E-6 ± 1.10E-6 | 8.46E-7 ± 3.14E-7 | 1.29E-7 ± 5.99E-8 | 2.39E-6 ± 4.89E-7 | 1.39E-7 ± 2.52E-8 | 1.01E-7 ± 2.64E-8 |
| Total | $1 \times 10^{-6}$ | 3 | 11/05/2021 | 10:35 a.m. | D7 | 1.19E-5 ± 2.45E-6 | 2.74E-6 ± 9.92E-7 | 1.05E-6 ± 4.22E-7 | 1.00E-7 ± 6.11E-8 | 3.44E-6 ± 1.77E-7 | 1.58E-7 ± 1.52E-8 | 1.45E-7 ± 3.08E-8 |
| Total | $5 \times 10^{-6}$ | 3 | 11/05/2021 | 10:40 a.m. | D7 | 4.21E-5 ± 2.79E-5 | 6.17E-6 ± 1.60E-6 | 1.38E-6 ± 7.36E-7 | 1.94E-7 ± 3.30E-8 | 2.32E-6 ± 1.77E-7 | 2.72E-7 ± 7.70E-8 | 2.00E-7 ± 9.47E-8 |
| Total | $1 \times 10^{-5}$ | 3 | 11/05/2021 | 10:45 a.m. | D7 | 3.74E-5 ± 1.07E-5 | 4.13E-6 ± 1.55E-6 | 1.47E-6 ± 4.60E-7 | 1.06E-7 ± 7.92E-8 | 2.69E-6 ± 1.14E-6 | 1.54E-7 ± 2.29E-8 | 1.37E-7 ± 3.56E-8 |

**Table S4.** Raw metabolomic data set.

.xlsx file

**Table S5.** PLS-DA cross-validation details.

| Measure | 1 comps | 2 comps | 3 comps | 4 comps | 5 comps |
|---|---|---|---|---|---|
| Accuracy | 0.088529 | 0.12279 | 0.18985 | 0.24662 | 0.29015 |
| R2 | 0.5242 | 0.66048 | 0.7778 | 0.85191 | 0.89565 |
| Q2 | 0.42734 | 0.5728 | 0.67829 | 0.75186 | 0.78209 |

**Table S6.** ECDF parameters.

.xlsx file

**Figures**

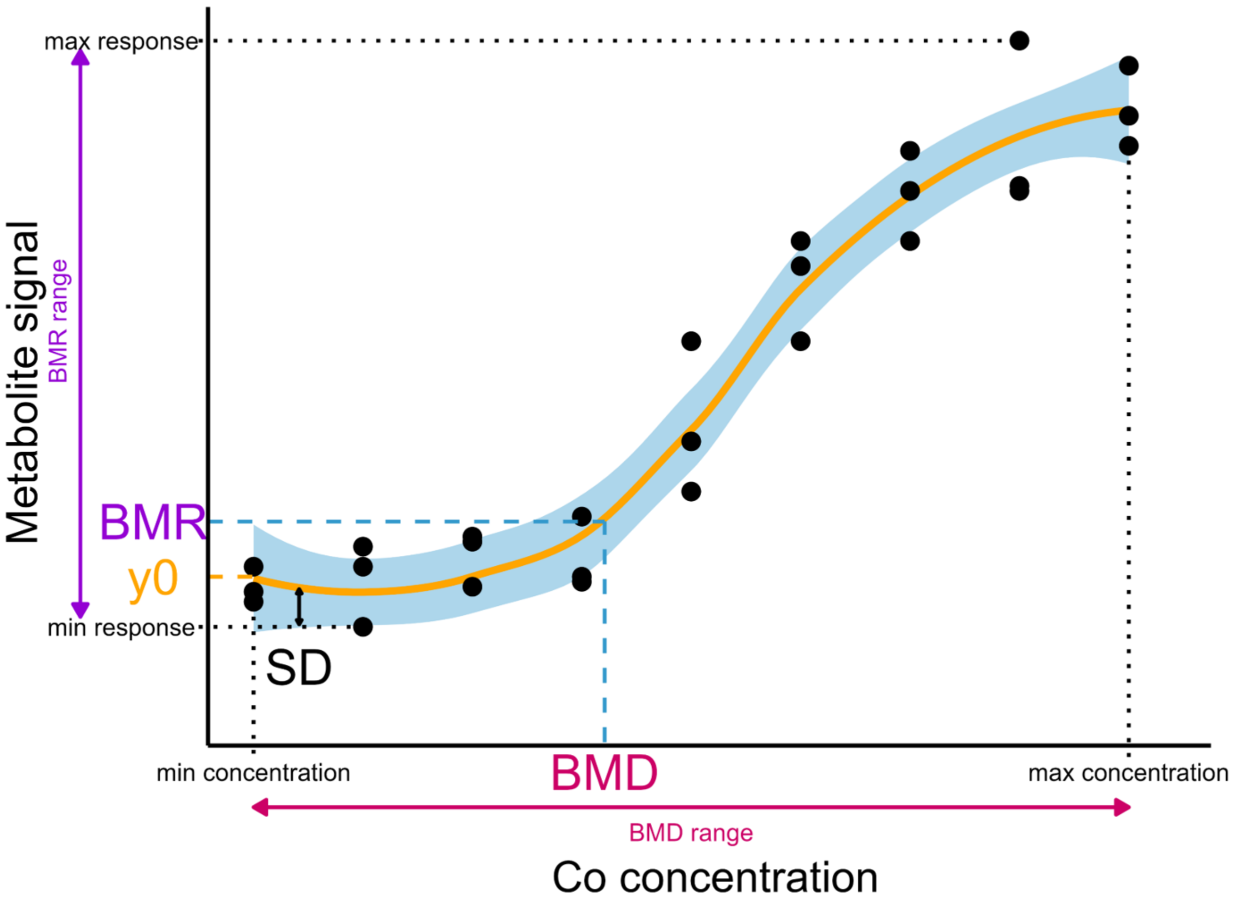


**Figure S1**: Benchmark-dose calculation illustration.

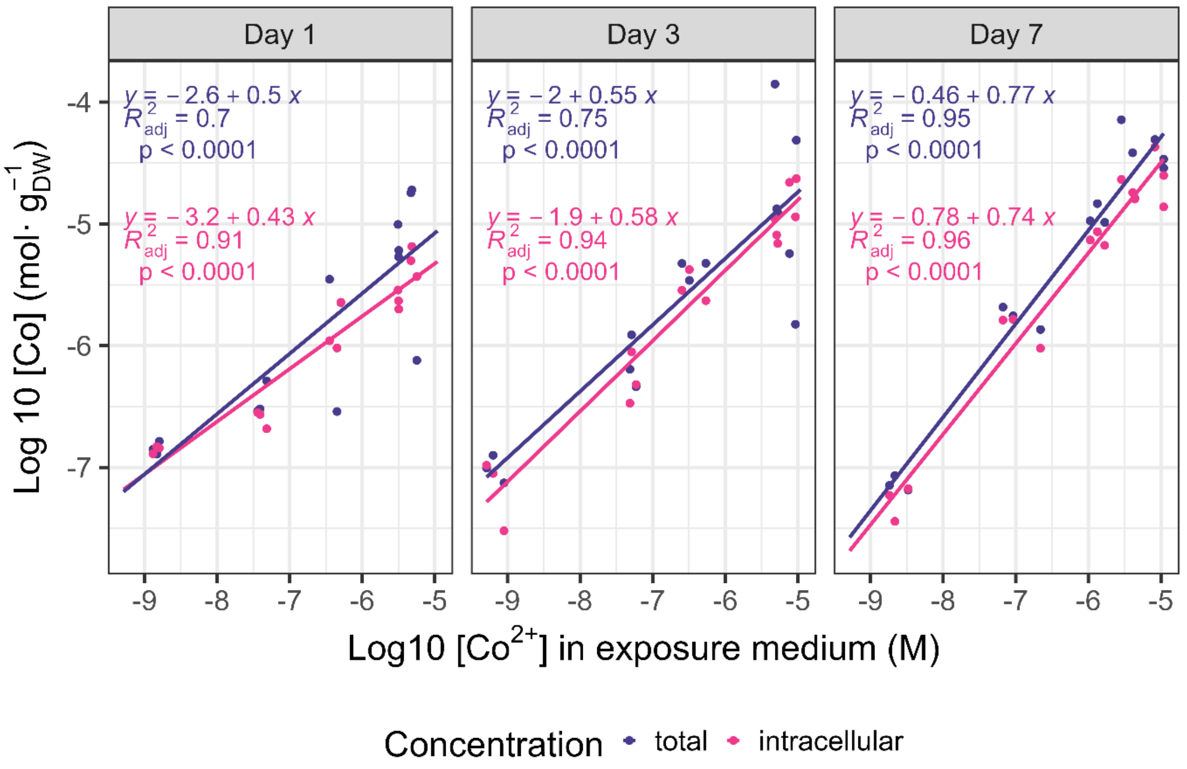


**Figure S2:** Cobalt bioaccumulation of biofilms. Total and intracellular Co concentrations ($\log_{10}$ mol·$g^{-1}_{DW}$) as function of $Co^{2+}$ concentration ($\log_{10}$ M) in exposure medium at different exposure time.

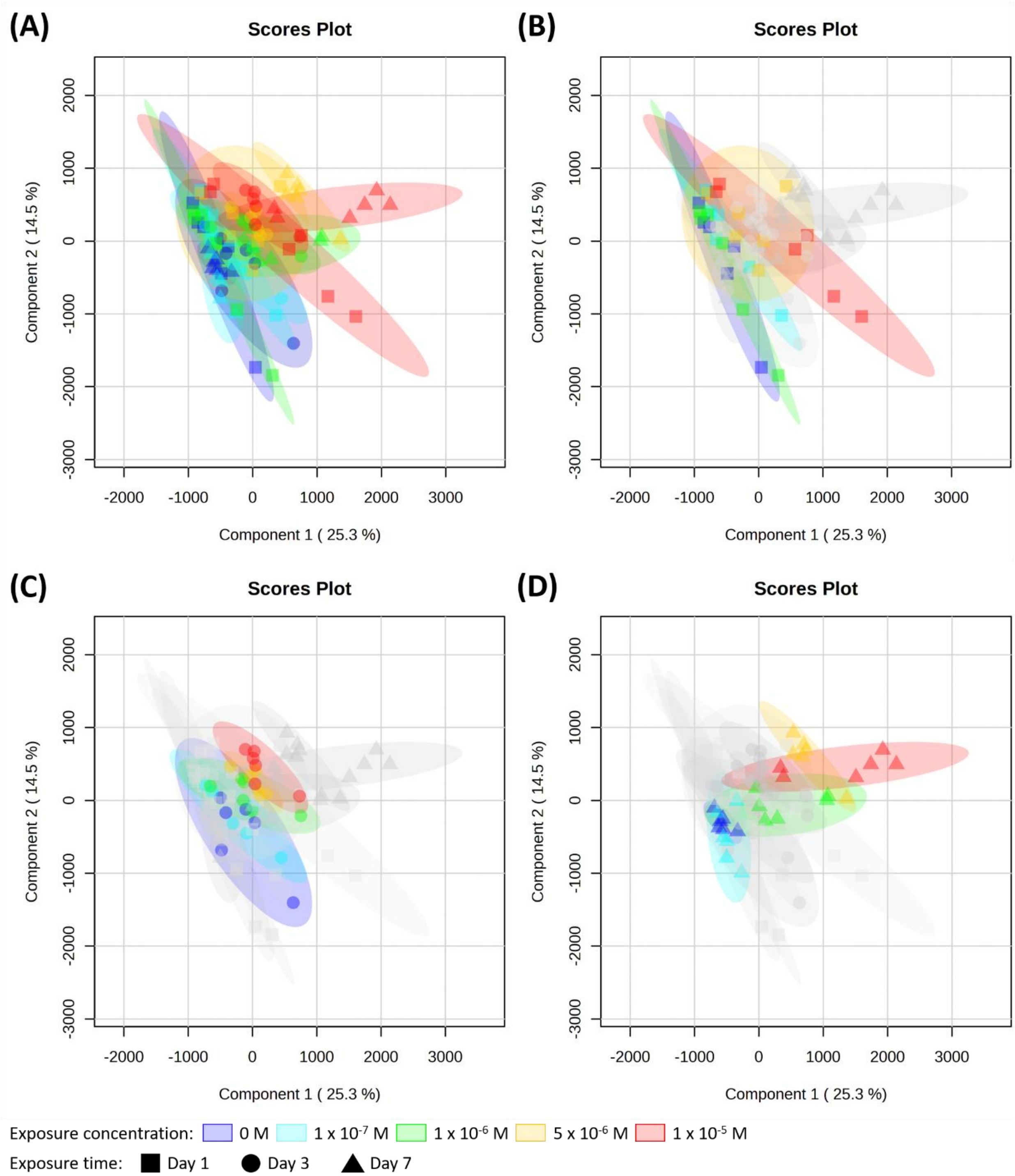

(A)
Scores Plot
(B)
Scores Plot
(C)
Scores Plot
(D)
Scores Plot
Component 2 ( 14.5 %)
Component 1 ( 25.3 %)
Exposure concentration: 0 M 1 x 10-7 M 1 x 10-6 M 5 x 10-6 M 1 x 10-5 M
Exposure time: Day 1 Day 3 Day 7

(E)

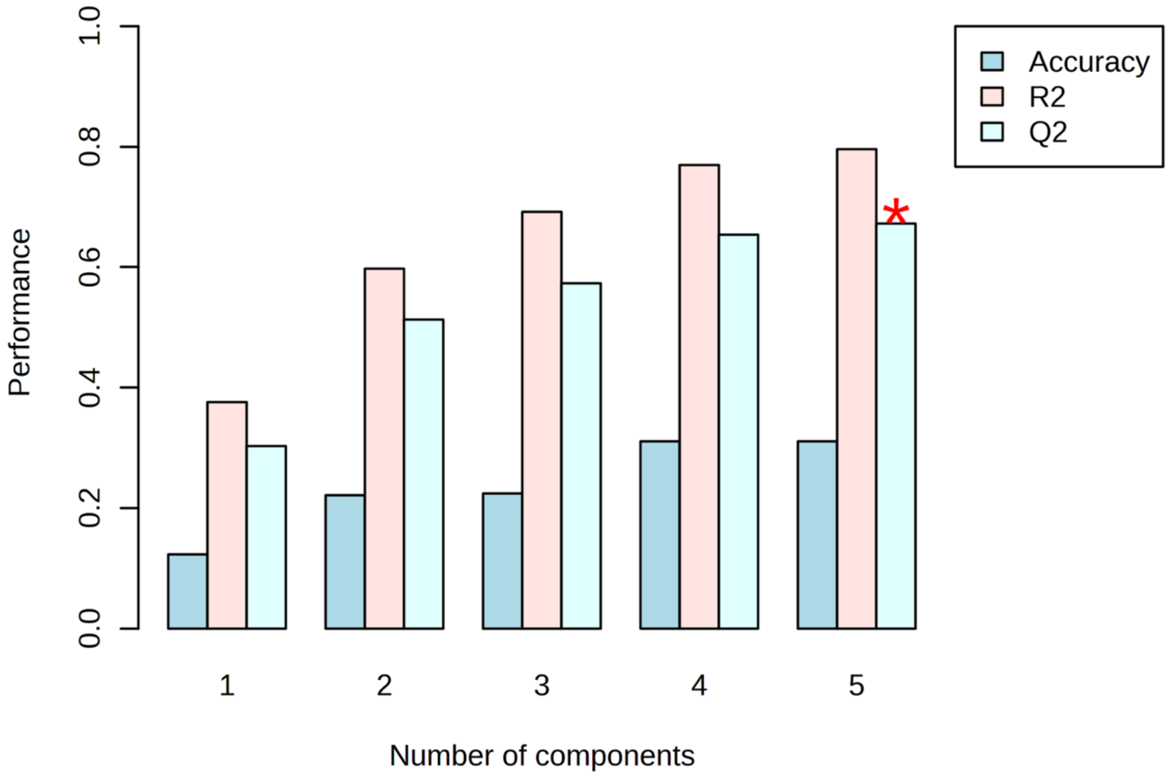

Accuracy
R2
Q2
Performance
0.0
0.2
0.4
0.6
0.8
1.0
1
2
3
4
5
Number of components


(F)

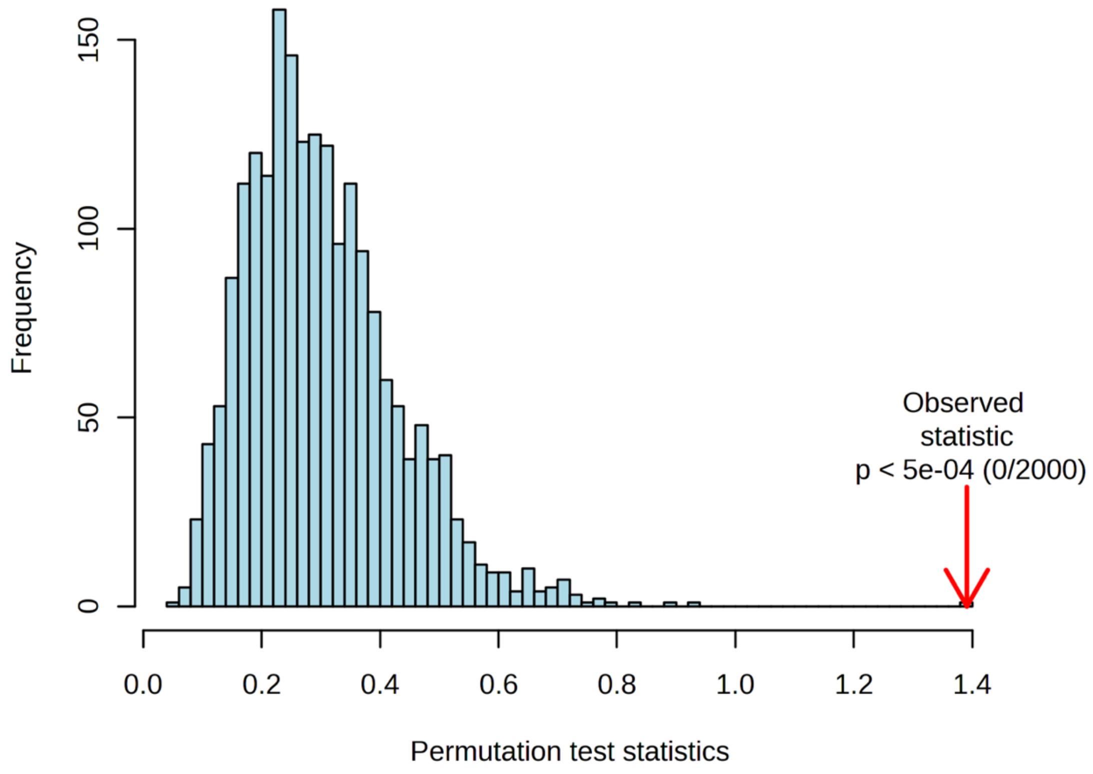

Frequency
0
50
100
150
Observed
statistic
p < 5e-04 (0/2000)
0.0
0.2
0.4
0.6
0.8
1.0
1.2
1.4
Permutation test statistics

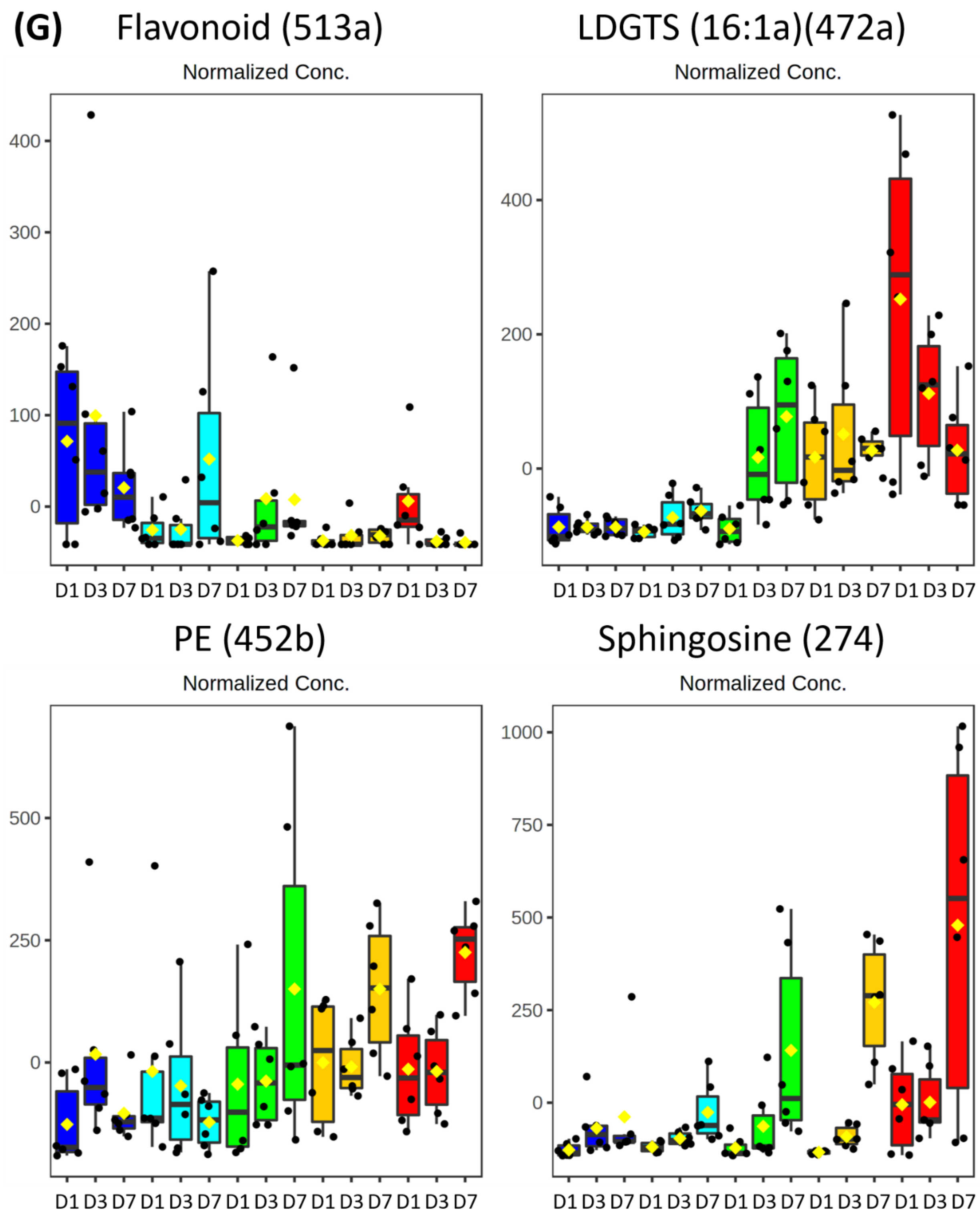

(G)
Flavonoid (513a)
LDGTS (16:1a)(472a)
PE (452b)
Sphingosine (274)
Normalized Conc.
D1 D3 D7 D1 D3 D7 D1 D3 D7 D1 D3 D7 D1 D3 D7

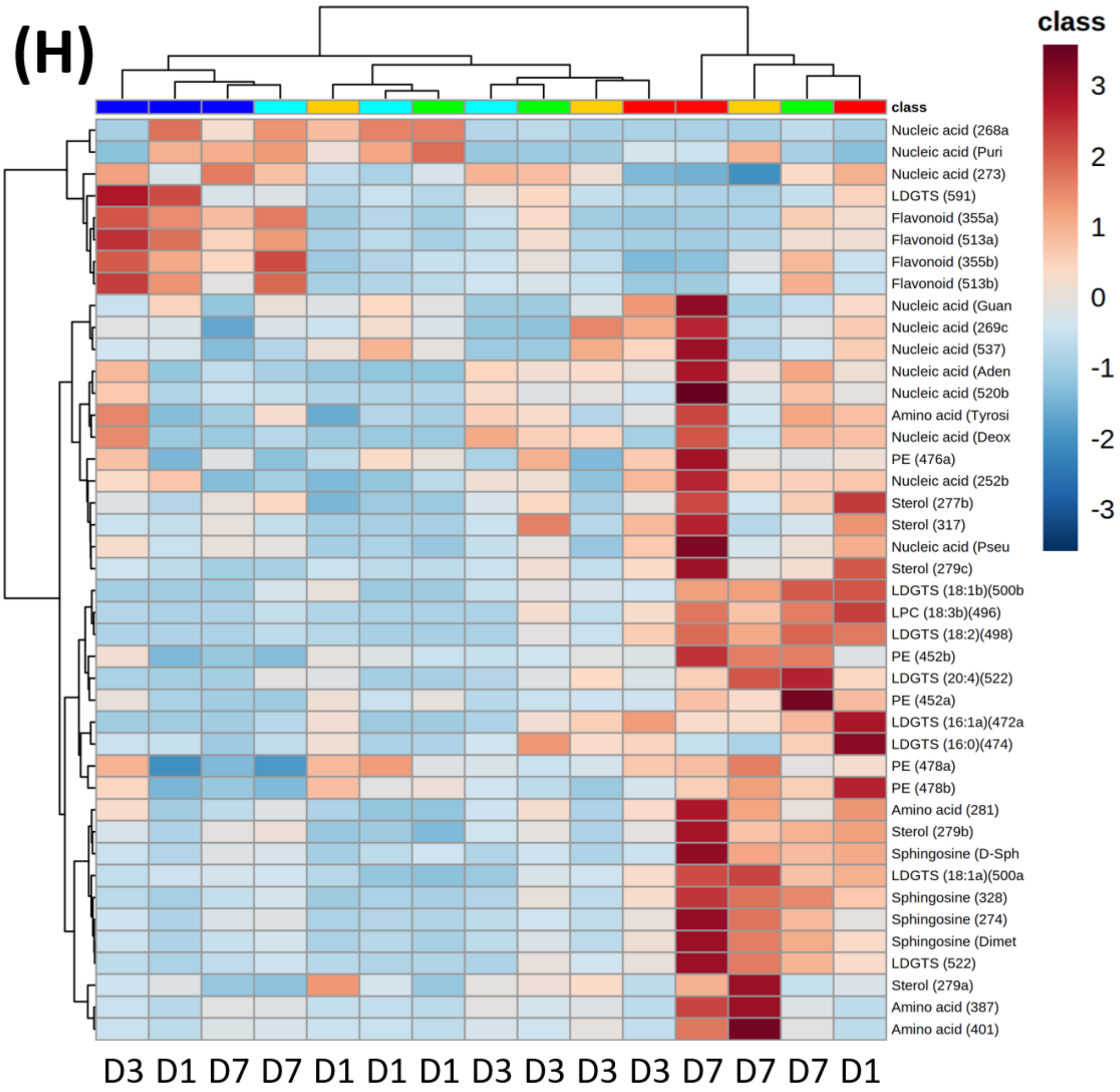


**Figure S3:** Meta-metabolomic response of biofilms according to time and concentrations of exposure to Co. (A) Individual score plot generated from PLS-DA analysis performed with 159 annotated variables to components 1-2 with all conditions grouped together, (B) after one day of exposure, (C) three and (D) seven. Quality descriptors of the statistical significance and predictive ability of the discriminant model according to (E) cross-validations test and (F) corresponding permutations. (G) Four examples of representative box-plot. (H) Heatmap with hierarchical classification representation of concentration class averages (Ward clustering according to Euclidian distances) performed from relative intensities of the dysregulated analytes with a VIP score > 1 for the component 1.

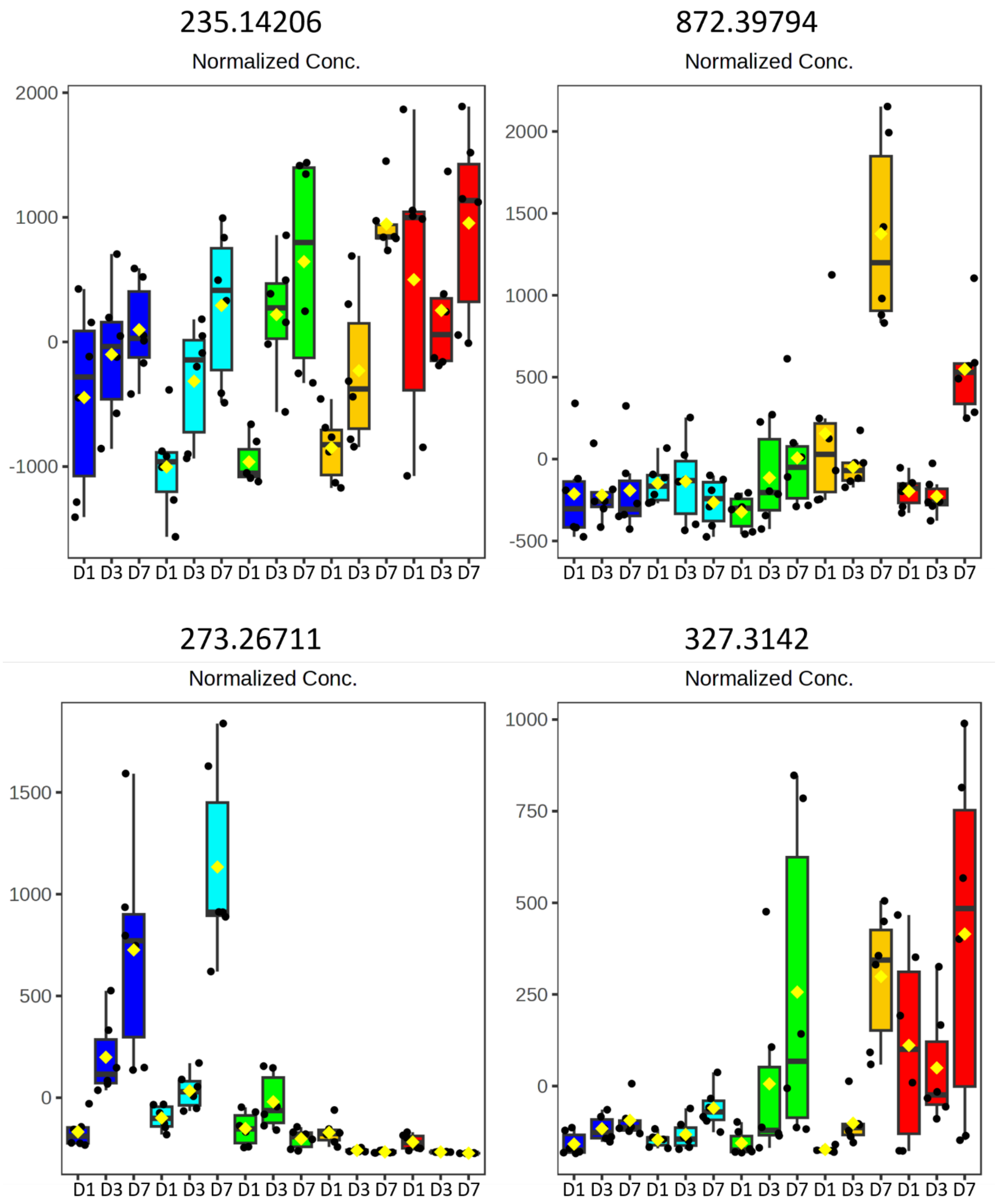


**Figure S4.** Four examples of representative box plots of metabolite fluctuations

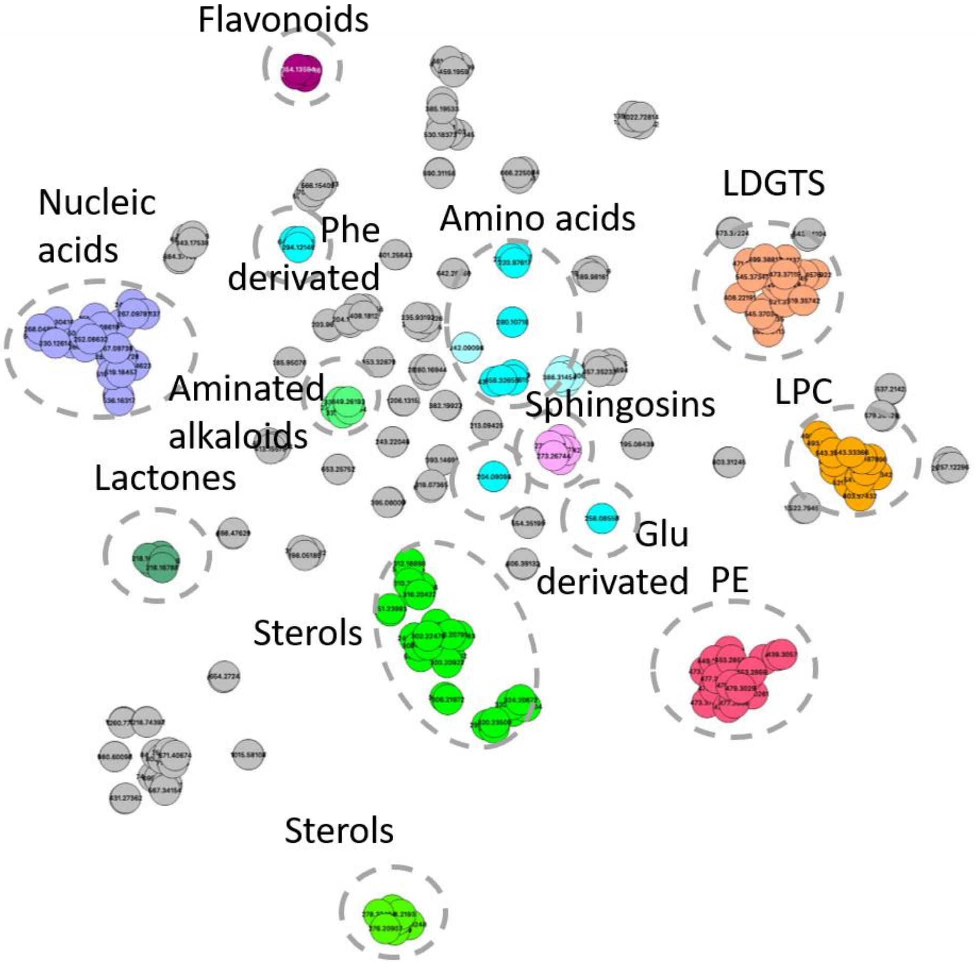


**Figure S5**: t-distributed Stochastic Neighbor Embedding of annotated metabolites.

REFERENCES

(1) Bryan, S. E.; Tipping, E.; Hamilton-Taylor, J. Comparison of Measured and Modelled Copper Binding by Natural Organic Matter in Freshwaters. *Comp. Biochem. Physiol. Part C Toxicol. Pharmacol.* **2002**, *133* (1–2), 37–49. https://doi.org/10.1016/S1532-0456(02)00083-2.

(2) Tipping, E. Modelling the Interactions of Hg(II) and Methylmercury with Humic Substances Using WHAM/Model VI. *Appl. Geochem.* **2007**, *22* (8), 1624–1635. https://doi.org/10.1016/j.apgeochem.2007.03.021.

(3) Jeffrey, S. W.; Humphrey, G. F. New Spectrophotometric Equations for Determining Chlorophylls a, b, C1 and C2 in Higher Plants, Algae and Natural Phytoplankton. *Biochem. Physiol. Pflanz.* **1975**, *167* (2), 191–194. https://doi.org/10.1016/S0015-3796(17)30778-3.
